# The extended language network: Language-responsive brain areas whose contributions to language remain to be discovered

**DOI:** 10.1101/2025.04.02.646835

**Authors:** Agata Wolna, Aaron Wright, Colton Casto, Samuel Hutchinson, Benjamin Lipkin, Evelina Fedorenko

## Abstract

Although language neuroscience has largely focused on ‘core’ left frontal and temporal brain areas and their right-hemisphere homotopes, numerous other areas—cortical and subcortical—have been implicated in linguistic processing. However, these areas’ contributions to language remain unclear given that the evidence for their recruitment comes from diverse paradigms, many of which conflate language processing with perceptual, motor, or task-related cognitive processes. Using fMRI data from 772 participants (438 females, 334 males) performing an extensively validated language ‘localizer’ paradigm that isolates language processing from other processes, we a) delineate a comprehensive set of areas that respond reliably to language across written and auditory modalities, and b) evaluate these areas’ selectivity for language relative to a demanding non-linguistic task. In line with prior claims, many areas outside the core fronto-temporal network respond during language processing, and most of them show selectivity for language relative to general task demands. These language-selective areas of the extended language network include areas around the temporal poles, in the medial frontal cortex, in the hippocampus, and in the cerebellum, among others. Although distributed across many parts of the brain, the extended language-selective network still only comprises a small fraction (<5%) of the grey matter volume, challenging the view that the entire brain processes language. These newly identified language-selective areas can now be systematically characterized to decipher their contributions to language processing, including testing whether these contributions differ from those of the core language areas.

**Significance statement:** Language processing consistently recruits a left-lateralized fronto-temporal brain network, but language tasks often additionally engage areas outside this core system. In an fMRI dataset of 772 participants performing a validated language localizer task, we identified 17 brain areas outside the core fronto-temporal network that respond to both auditory and written language. Most of these areas show selectivity for language, including regions in the temporal poles, medial frontal cortex, hippocampus, and cerebellum. Despite its large number of components, this extended language network still only takes up a small fraction of the grey matter volume, challenging the view that the entire brain processes language. These findings lay the foundation for systematic characterization of these newly identified non-canonical language areas.

## Introduction

Language processing engages a left-lateralized fronto-temporal brain network—the “language network“—which selectively supports comprehension and production across modalities, including computations related to word retrieval and combinatorial syntactic and semantic processing (Vigneau et al., 2006; Turker et al., 2023; Fedorenko et al., 2024). These areas have been consistently identified across diverse neuroimaging approaches (PET, fMRI, MEG, and intracranial recordings) and paradigms, and also implicated in studies of patients with aphasia (see **Table OSF1** for select reviews, meta-analyses, and empirical papers). In addition to this ‘core’ network and its right-hemispheric homotopic regions (Lindell, 2006; Ferstl et al., 2008; Mahowald & Fedorenko, 2016; Martin et al., 2022), several other brain areas have been implicated in aspects of language. In the cortex, these include medial frontal areas (Ardila, 2020; Bohsali & Crosson, 2016; Hertrich et al., 2016), medial and lateral parietal areas (Hasson et al., 2007; Peschke et al., 2012; Papagno et al., 2017), temporal poles (Ferstl et al., 2008; Holland & Lambon Ralph, 2010; Lambon Ralph et al., 2017), ventral temporal areas (Krauss et al., 1996; Bédos Ulvin et al., 2017; Snyder et al., 2023), and even occipital areas in the blind (Bedny et al., 2011; Reich et al., 2011; Kim et al., 2017) but also sighted populations (Dikker et al., 2009, 2010; Seydell-Greenwald et al., 2023). Subcortically, language-implicated areas include the hippocampus (Covington & Duff, 2016; Dijksterhuis et al., 2024; reviews: Duff & Brown-Schmidt, 2012, 2017; Piai et al., 2016), the thalamus (Wahl et al., 2008; review: Klostermann & Ehlen, 2013), and basal ganglia structures, including the caudate, the pallidum, and the putamen (Booth et al., 2007; Crosson et al., 2003; Kotz, 2009; Oberhuber et al., 2013; Thibault et al., 2021, 2025). Finally, parts of the cerebellum have long been implicated in speech and language (reviews: Murdoch, 2010; Mariën et al., 2014; LeBel & D’Mello, 2023).

However, the precise contributions of these non-canonical brain areas to language remain unclear. Much of the evidence comes from studies relying on single experimental paradigms, which often fail to isolate language processing from perceptual, motor, or general cognitive demands (for discussion, see Fedorenko et al., 2024). Further, the group-averaging approach—adopted by the majority of past neuroimaging studies—complicates across-study comparisons. In this approach, individual maps are averaged voxel-wise, and group-level activation peaks are reported in a standardized coordinate system (e.g., MNI; Mazziotta et al., 2001). However, inter-individual neuroanatomical variability leads to differences in structural and functional topographies (Frost & Goebel, 2012; Tahmasebi et al., 2012; Fedorenko, 2021; Michon et al., 2022), which may shift the group peaks across studies even for the same paradigm. Besides, any macrocoanatomical structure encompasses multiple functionally distinct areas, making it difficult to unambiguously determine whether nearby peaks reported in different studies correspond to the same functional area (earlier discussions: Brett et al., 2002; Saxe et al., 2006).

One approach that circumvents these challenges is functional localization (Saxe et al., 2006; Kanwisher, 2010; Fedorenko et al., 2010). Instead of averaging brains together, this approach identifies functional areas within individuals using ‘localizer’ tasks—robust contrasts that target a particular perceptual or cognitive process. This approach ensures that functionally equivalent area(s) are referred to across individuals and studies, supporting the systematic characterization of each area’s responses across paradigms, as needed to understand its computations. At the same time, identifying these areas and testing their properties in each brain individually—effectively, working in each individual’s functional native space (regardless of the anatomical, stereotactic space that the data are registered to)—allows to preserve high functional resolution (Nieto-Castañón & Fedorenko, 2012): i.e., the ability to differentiate the relevant areas from nearby, functionally distinct areas.

Here, we use an established language localizer and a large fMRI dataset to search for language-selective areas outside the core language network. In line with past claims, we find several such areas, and many of them show selectivity for language relative to a demanding non-linguistic task, which makes them exciting targets for future investigations.

## Methods

### Overview

We leverage a dataset of 772 participants each of whom performed an extensively validated reading-based language localizer (Fedorenko et al., 2010) and one or two additional tasks (**Figure 1A**). In particular, 489 of the 772 participants performed a demanding spatial working memory task, which allowed us to begin evaluating the selectivity of the non-canonical language-responsive areas. 86 of these 489 participants additionally performed an auditory version of the language localizer (Malik-Moraleda, Ayyash et al., 2022), which allowed us to evaluate whether the observed language responses are robust to modality (reading vs. listening). If a brain region, akin to the core frontal and temporal language areas, responds to both written and auditory linguistic inputs (Fedorenko et al., 2010; Vagharchakian et al., 2012; Regev et al., 2013; Deniz et al., 2019), its computations plausibly have to do with the content of the linguistic signals rather than their surface (visual or auditory) properties.

**Figure 1.**
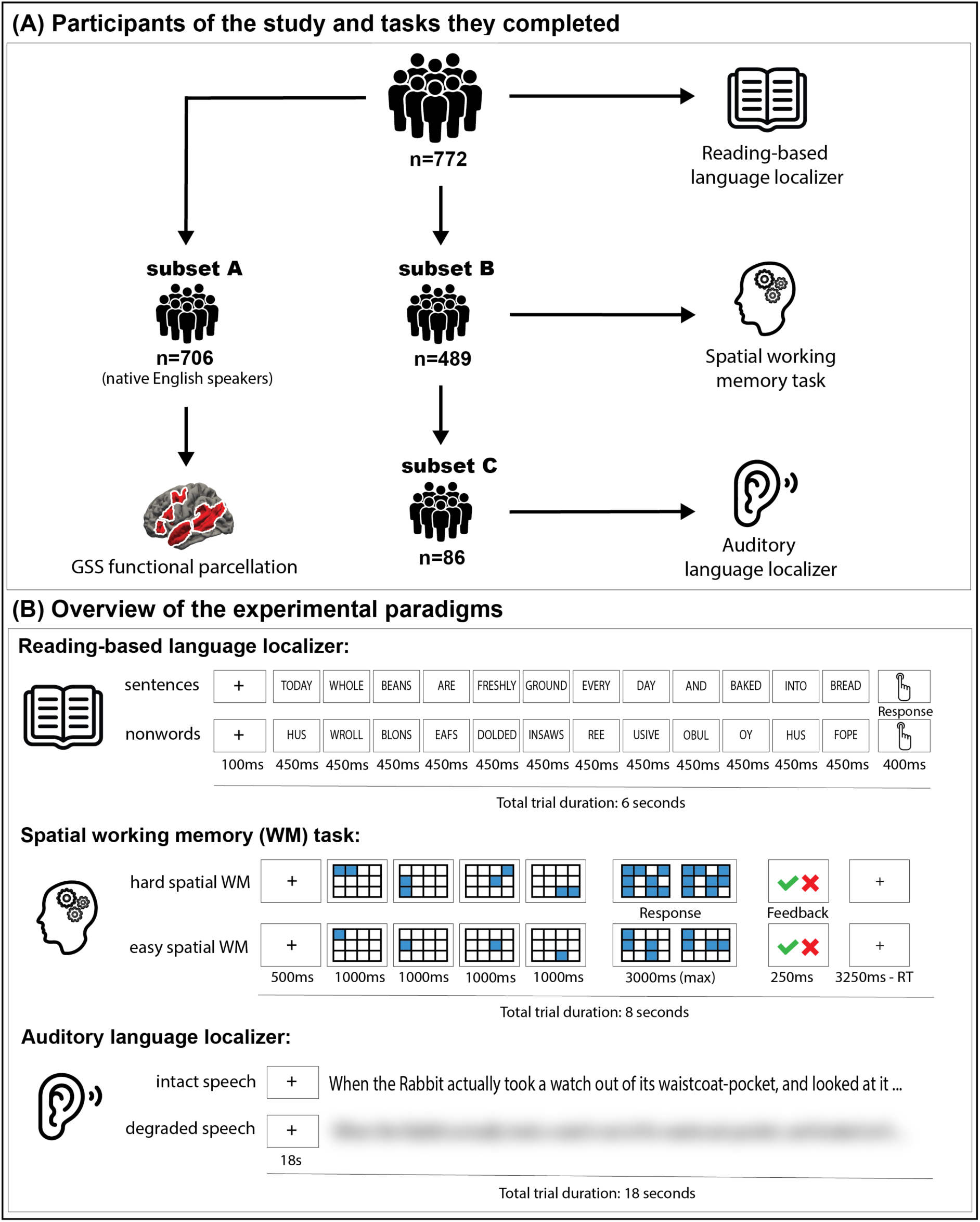
Overview of the dataset and paradigms. A) The structure of the dataset. A total of n=772 participants were included. All 772 completed the reading-based language localizer. A subset of n=706 (*subset A*) were native speakers of English and were used in the GSS analysis for creating the extended set of parcels. A subset of n=489 participants (*subset B*) completed the spatial working memory task, and a subset of n=86 participants (*subset C*) completed the auditory language localizer. B) The paradigms. For each task, a sample trial in each condition is shown (see <u>Tasks</u> for a detailed description).

In the critical analysis, we define regions of interest functionally within individual participants. To constrain the selection of these individual-level functional regions of interest (fROIs), we use functional parcels derived from a group-level representation of the language localizer activity (Fedorenko et al., 2010). Because the fMRI signal is weaker in subcortical areas, which are further away from the receiver coil, and to ensure that we don’t miss any language-responsive areas, we complement these parcels with subcortical anatomical parcels, which we use in the same way (to constrain individual-level fROIs). Finally, because standard cortical atlases are commonly used for ROI definition, we comprehensively examine language responsiveness and selectivity in three such atlases, in order to specify the relevant subsets of areas within which reliable and selective responses to language can be observed, and to further highlight the importance of functional localization within individuals instead of using group-level anatomical masks.

### Participants

The structure of the dataset is shown in **Figure 1A**. We used three tasks (**Figure 1B**): a standard reading-based language localizer (Fedorenko et al., 2010), an auditory language localizer based on excerpts from *Alice in Wonderland* (Malik-Moraleda, Ayyash et al., 2022), and a spatial working memory task (Fedorenko et al., 2013; Assem et al., 2020).

772 neurotypical adults (age: mean=27, range=18-81; 438 females) each completed the standard reading-based language localizer (Fedorenko et al., 2010). A subset of 706 participants (age: mean=27, range=18-81; 404 females; *subset A*), all native speakers of English, were included in the GSS (Group-constrained Subject-Specific) analysis (Fedorenko et al., 2010), to create a set of extended language parcels (<u>Group-level parcels used to constrain the definition of individual fROIs</u>). 489 of the 772 participants (age: mean=29, range=17-80; 225 females; *subset B*) performed a spatial working memory task (Fedorenko et al., 2013; Assem et al., 2020), and 86 of these 489 participants (age: mean=28, range=19-45; 44 females; *subset C*) additionally performed an auditory language localizer task. This subset consisted of 20 native English speakers and 66 native speakers of diverse languages (all proficient in English), who performed the auditory localizer in their native language (data partially reported in (Malik-Moraleda, Ayyash et al., 2022). All participants gave written informed consent, in accordance with the requirements of MIT’s Committee on the Use of Humans as Experimental Subjects (COUHES) and were paid for their participation.

### Tasks

#### Reading-based language localizer task

Participants passively read English sentences and lists of pronounceable nonwords, which appeared on the screen one word/nonword at a time (**Figure 1B, top**). Each trial started with a 100ms fixation, followed by a 12-element-long sequence of words/nonwords presented at the rate of 450ms. At the end of each trial, a drawing of a hand appeared for 400ms, and participants had to press a button every time they saw this drawing. The button press task was included to help the participants stay alert and focused during the task. Each trial ended with a blank screen presented for 100ms. The task followed a blocked design. The total duration of each trial was 6 seconds, and 3 trials were included in each block. Each run included 16 18-second blocks (8 per condition). Additionally, 5 fixation blocks of 14 seconds were included in each run, for a total run duration of 358 seconds. Each participant completed two runs. The *Sentences* > *Nonwords* contrast targets brain areas that support language processing (including mental operations related to word retrieval, syntactic structure building, and semantic composition), while excluding perceptual processing (see **Figure OSF1**—available at https://osf.io/7594t/—for evidence that this contrast includes areas sensitive to sub-lexical/phonological processing; also: Bozic et al., 2015; Regev et al., 2022).

It is important to note that although this particular version of the language localizer uses written stimuli, the core frontal and temporal areas activated by the written *Sentences>Nonwords* contrast overlap almost perfectly with areas identified with auditory contrasts between language processing and a perceptually similar condition, such as backwards speech or acoustically degraded speech (e.g., Fedorenko et al., 2010; Scott et al., 2017; Malik-Moraleda, Ayyash et al., 2022). Moreover, the network of areas that emerges for the *Sentences>Nonwords* contrast (and similar contrasts) closely matches one of the intrinsic brain networks recoverable from individual patterns of functional connectivity, as measured during naturalistic conditions such as resting state (e.g., Braga et al., 2020; Du et al., 2024) or even during task paradigms (e.g., Du et al., 2025; Shain & Fedorenko, 2025).

#### Auditory language localizer task

Participants passively listened to intact and acoustically degraded passages from *Alice in Wonderland* (Carroll, 1865; **Figure 1B, middle**). The degraded versions of each passage were created using a procedure introduced in Scott et al. (2017): intact passages were low-pass filtered (passband 500 Hz) and mixed with a time-varying noise track derived from the same clip (generated by randomizing consecutive 0.02-s segments, amplitude-modulated by the intact signal, and low-pass filtered to attenuate very high frequencies). The noise level was adjusted until the resulting clips were unintelligible. Although the degraded clips largely preserve prosodic patterns, the key computations carried out by the language areas (related to word recognition, syntactic structure building, and semantic composition; e.g., Shain, Kean et al., 2024) cannot be performed because the content is not comprehensible, leading to a much weaker response compared to the response to the intact clips.

Each trial consisted of an 18-second recording. Each run included 12 18-second trials (4 per condition; in addition to the intact and degraded conditions, the experiment included a third condition not included in any analyses here—recordings in an unfamiliar foreign language). Additionally, 3 fixation blocks of 12 seconds were included in each run, for a total run duration of 252 seconds. Each participant completed three runs. Similar to the reading-based localizer, the *Intact* > *Degraded* contrast targets brain areas that support language processing.

#### A demanding non-linguistic (spatial working memory) task

Participants performed a spatial working memory task consisting of easier and harder trials (e.g., Fedorenko et al., 2013; Assem et al., 2020). Each trial started with a 500ms fixation. Then, a 3 x 4 grid appeared in the center of the screen, and a series of locations were flashed in blue at the rate of 1 second per flash (**Figure 1B, bottom**). In the easy condition, one location flashed at a time (four locations in total); in the hard condition, two locations flashed at a time (eight in total). Participants were instructed to keep track of the locations. Each trial ended with a two-choice forced choice task: two grids with different sets of locations were shown side by side, and participants had to choose the grid that shows the locations just presented. They were told whether they chose correctly (green checkmark) or not (red cross). The task followed a blocked design. The total duration of each trial was 8 seconds, and 4 trials were included in each block. Each run included 12 32-second blocks (6 per condition). Additionally, 4 fixation blocks of 16 seconds were included in each run, for a total run duration of 448 seconds. Each participant completed 2 runs. The *Hard* > *Easy* contrast targets brain areas that support demanding cognitive tasks, including mental operations associated with working memory, attention, and cognitive control.

### fMRI data acquisition

Structural and functional data collected in the Athinoula A. Martinos Imaging Center at the McGovern Institute for Brain Research at MIT. Structural and functional data were acquired using a Siemens Trio 3 Tesla scanner before or after a Prisma upgrade (n=508 and n=264, respectively) and a 32-channel head coil. High-resolution, whole-brain anatomical images were acquired using two different T1-weighted MPRAGE sequence (**Sequence 1**: n=508, 176 sagittal slices; voxel size 1 x 1 x 1 mm^3^; TR = 2530 ms, TE = 3.48 ms, flip angle = 9°; **Sequence 2**: n=264, 208 sagittal slices; voxel size 0.9 x 0.9 x 0.9 mm^3^; TR = 1800 ms, TE = 2.40 ms, flip angle = 9°). Functional T2*-weighted images were acquired using three different whole-brain echo-planar (EPI) pulse sequences with phase encoding direction A >> P (**Sequence 1, Siemens Trio**: n = 508, 31 near-axial slices, 2.1 × 2.1 × 4 mm voxels; TR = 2,000 ms; TE = 30 ms; flip angle = 90°; matrix size 96 x 96 mm; MB acceleration factor 0; **Sequence 2, Siemens Prisma**: n = 73, 52 near-axial slices, 2 × 2 × 2 mm isotropic voxels; TR = 2,000 ms; TE = 30 ms; flip angle = 90°; matrix size 104 × 104 mm; MB acceleration factor 2; **Sequence 3, Siemens Prisma**: n = 191, 72 near-axial slices, 2 × 2 × 2 mm isotropic voxels; TR = 2,000 ms; TE = 30 ms; flip angle = 90°; matrix size 104 × 104 mm; MB acceleration factor 3). The first 10 s of each run were excluded to allow for steady state magnetization.

### fMRI data preprocessing

fMRI data were analyzed using SPM12 (release 7487), CONN EvLab module (release 19b), and other custom MATLAB scripts. Each participant’s functional and structural data were converted from DICOM to NIFTI format. All functional scans were coregistered and resampled using 4^th^ degree B-spline interpolation to the first scan of the first session (Friston et al., 1995). Potential outlier scans were identified from the resulting subject-motion estimates as well as from BOLD signal indicators using default thresholds in CONN preprocessing pipeline (5 standard deviations above the mean in global BOLD signal change, or framewise displacement values above 0.9 mm; Nieto-Castañón, 2020). Functional and structural data were independently normalized into a common space (the Montreal Neurological Institute [MNI] template; IXI549Space) using SPM12 unified segmentation and normalization procedure (Ashburner & Friston, 2005) with a reference functional image computed as the mean functional data after realignment across all timepoints omitting outlier scans. The output data were resampled to a common bounding box between MNI-space coordinates (−90, −126, −72) and (90, 90, 108), using 2mm isotropic voxels and 4^th^ degree B-spline interpolation for the functional data, and 1mm isotropic voxels and trilinear interpolation for the structural data. Last, the functional data were smoothed spatially using spatial convolution with a 4 mm FWHM Gaussian kernel (following a reviewer’s request, we additionally provide unsmoothed activation maps on OSF (https://osf.io/7594t/); see also **Figure OSF2** for evidence that individual-level activation maps are highly similar regardless of the preprocessing approach: with vs. without normalization to the common space, and between our volumetric SPM-based pipeline and a surface-based Freesurfer pipeline).

### fMRI data first-level modeling

Responses in individual voxels were estimated using a General Linear Model (GLM) in which each experimental condition was modeled with a boxcar function convolved with the canonical hemodynamic response function (HRF) (fixation was modeled implicitly). Temporal autocorrelations in the BOLD signal timeseries were accounted for by a combination of high-pass filtering with a 128 seconds cutoff, and whitening using an AR(0.2) model (first-order autoregressive model linearized around the coefficient a=0.2) to approximate the observed covariance of the functional data in the context of Restricted Maximum Likelihood estimation (ReML). In addition to experimental condition effects, the GLM design included first-order temporal derivatives for each condition (included to model variability in the HRF delays), as well as nuisance regressors to control for the effect of slow linear drifts, subject-specific motion parameters (6 parameters), and potential outlier scans (identified during preprocessing as described above) on the BOLD signal.

### Group-level parcels used to constrain the definition of individual fROIs

The group-level functional parcels were created using the group-constrained subject-specific approach (GSS; Fedorenko et al., 2010) as implemented in the spm_ss toolbox (available for download from: https://www.evlab.mit.edu/resources). This approach helps determine which parts of the activation landscape are consistent across participants and establish correspondences across individuals. To do so, thresholded binarized individual maps for a particular contrast of interest are overlaid in a common brain space to create a probabilistic activation overlap map (or ‘atlas’). Subsequently, this map is divided into parcels using a watershed image parcellation algorithm (see Julian et al., 2012 for the application of this approach to the ventral visual areas).

Here, we used a set of 706 individual maps (the subset of the native English speakers; **Figure 1A**) and selected the 20% most language-responsive voxels across the brain in each individual. This relatively liberal individual-level threshold increases the likelihood of capturing all language-responsive areas, including those with weaker and less spatially consistent responses. It is worth noting that the average *t-*value across participants for these voxels was still quite high: 2.02 (SD=0.6), with the average minimum *t-*value of 0.92, and the average maximum *t*-value of 14.17). In the resulting overlap map, the value of each voxel, divided by the total number of participants, represents the proportion of participants for whom that voxel belongs to the top 20% of most language-responsive voxels (the highest value in this map is ∼0.78, in line with Lipkin et al., 2022). The overlap map was thresholded to remove voxels with values below 0.1 (i.e., voxels present in fewer than 10%, or 71 of the 706 participants) and smoothed with a 6 mm full-width half-maximum Gaussian kernel, to avoid over-parcellation. Finally, an image parcellation (watershed) algorithm was run to identify the main “hills” in the activation landscape (Meyer, 1991).

In our dataset, this algorithm identified 27 parcels (of size 150 voxels or larger): 11 in the left hemisphere, 8 in the right hemisphere, 3 midline bilateral parcels, and 5 parcels in the cerebellum (**Figures 2** and **3** and **Table 1**; see **Figure OSF3** for evidence that the probabilistic overlap map and the parcels are similar if the individual maps are thresholded more conservatively—at top 10% of most language-responsive voxels instead of top 20%). Six additional parcels (smaller than 150 voxels in size) were excluded from the main analysis, but we report their profiles for completeness in **Figure S1**.

**Figure 2.**
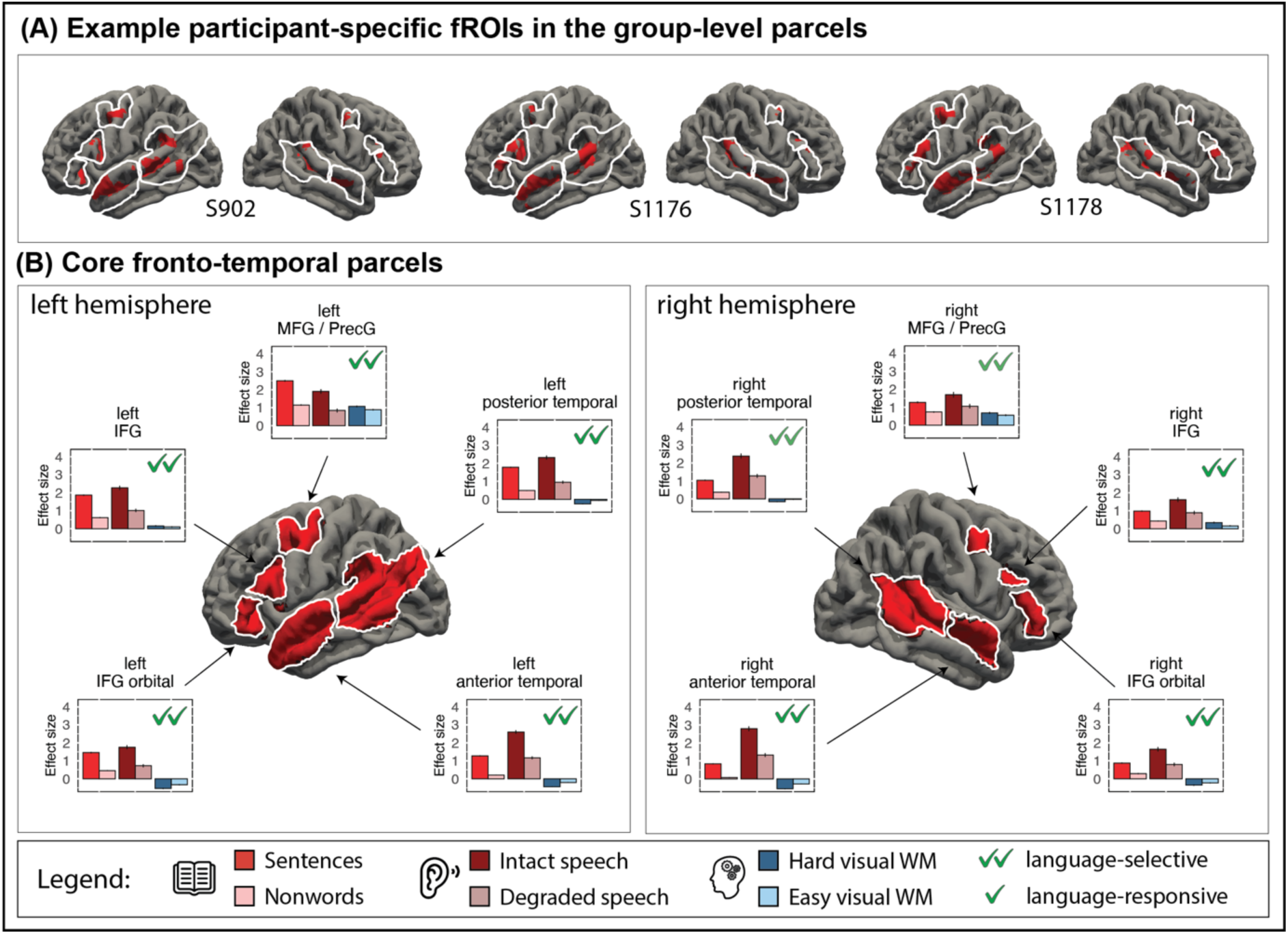
Language-responsive regions in the core language network and their functional profiles. A) Example participant-specific fROIs created using top 10% of most active voxels (shown in red) within the group-level parcels (white outlines) for the core language regions. B) The five core left-hemisphere parcels (left sub-panel) and their right-hemisphere homotopes (right sub-panel). The figure shows the surface projections of the parcels (all the analyses were performed in the volume space, as described in <u>Methods</u>; the surface projections were created using FreeSurfer). For each region (defined based on individual activation maps, as described in <u>Methods</u> and illustrated in panel A), we show responses to the reading-based and auditory language localizers (red bars; darker bars correspond to the critical conditions) and the non-linguistic spatial working memory task (blue bars; the darker bar corresponds to the harder condition). For the reading-based localizer, the responses were estimated using independent runs of the data. Parcels that contain language-responsive fROIs have a green checkmark in the upper right-hand corner; the ones where the fROIs are additionally language-selective have two green checkmarks (all fROIs in this set are language-selective). In all plots, the bars correspond to the mean of raw data and whiskers represent SEM of raw means.

**Figure 3.**
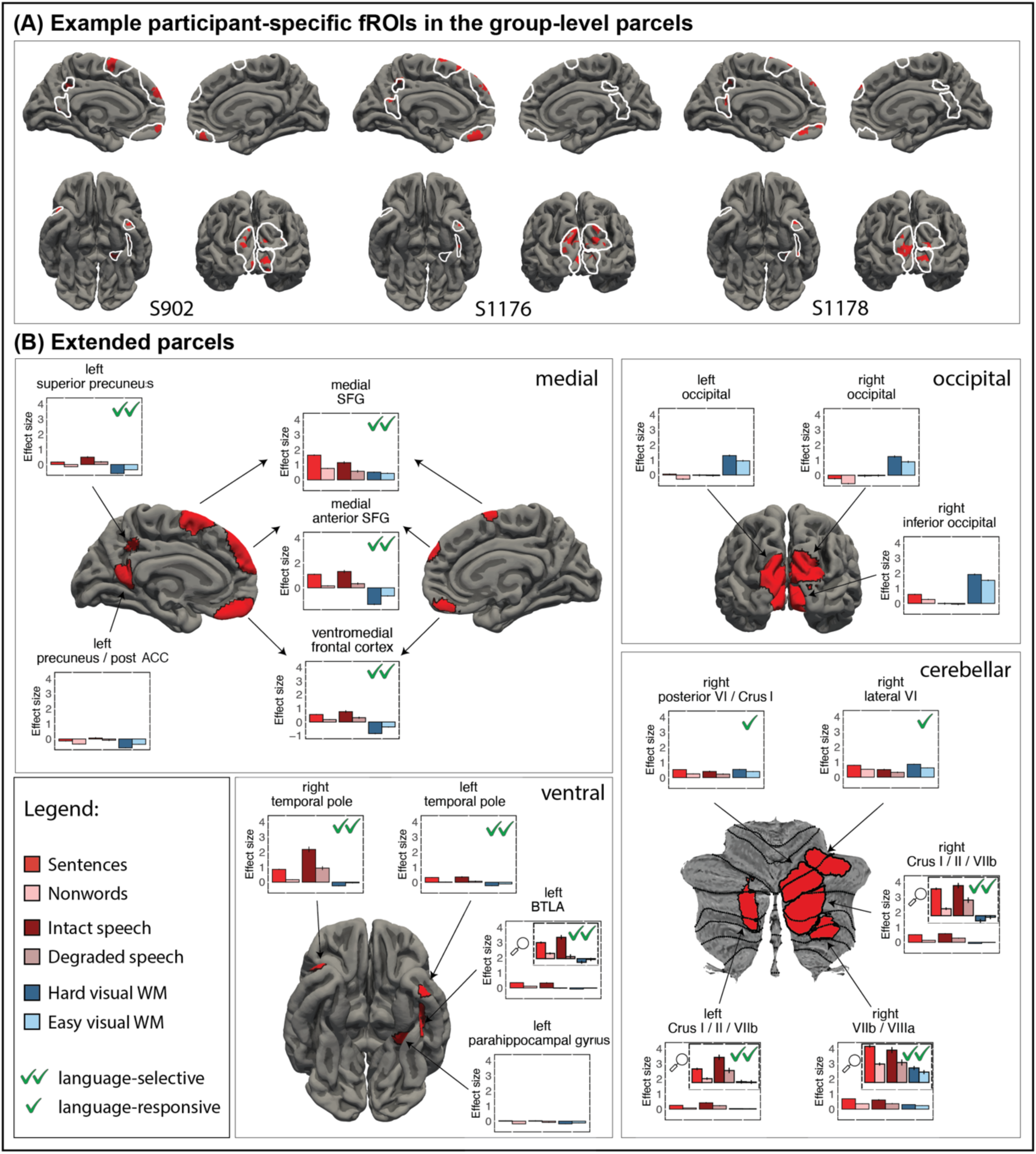
Language-responsive regions in the extended language network and their functional profiles. A) Example participant-specific fROIs created using top 10% of most active voxels (shown in red) within the group-level parcels (white outlines) for the non-canonical language regions. Note that due to image orientation in some cases, the fROIs are not visible in the surface projection (statistical maps for all subjects are available on OSF: https://osf.io/7594t/). B) The extended set of parcels (as identified by the GSS procedure; <u>Methods</u>). Separate sub-panels show the medial, occipital, ventral, and cerebellar parcels. The figure shows the surface projections of the cortical and cerebellar parcels (all the analyses were performed in the volume space, as described in <u>Methods</u>; the surface projections were created using FreeSurfer for the cortical and SUIT for the cerebellar parcels and are only used for visualization). For each region (defined based on individual activation maps, as described in <u>Methods</u> and illustrated in panel A), we show responses to the reading-based and auditory language localizers (red bars; darker bars correspond to the critical conditions) and the non-linguistic spatial working memory task (blue bars; the darker bar corresponds to the harder condition). For the reading-based localizer, the responses were estimated using independent runs of the data. Parcels that contain language-responsive fROIs have a green checkmark in the upper right-hand corner (n=12 of the 17 fROIs satisfy this criterion); the ones where the fROIs are additionally language-selective have two green checkmarks (n=10 fROIs). We used the same range of values for the y-axis across all regions to highlight differences in response magnitudes, but for the language-selective regions with low responses, we include insets, marked with a magnifying glass, where we restrict the y-axis to make the profiles easier to see.

**Table 1.**
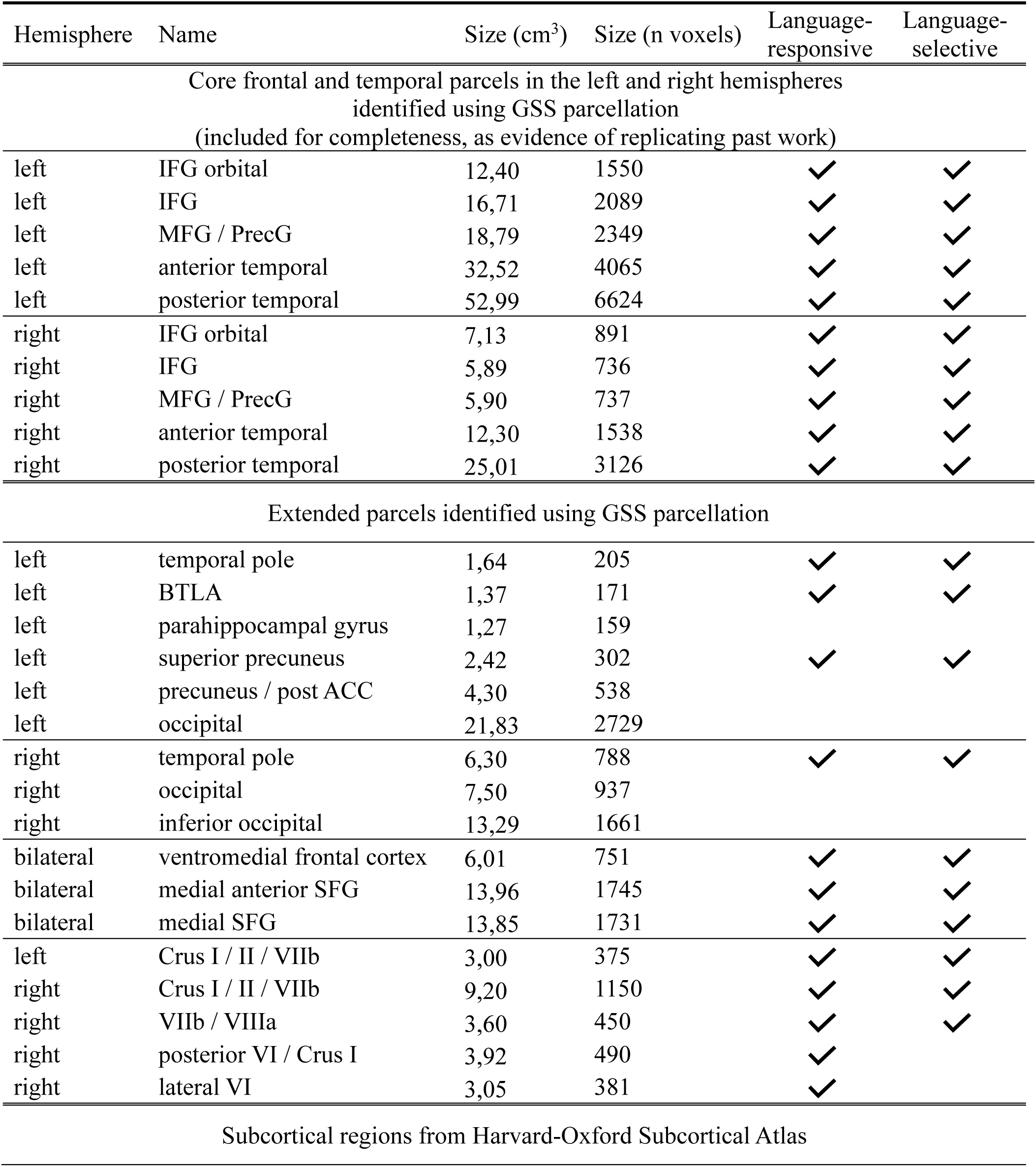
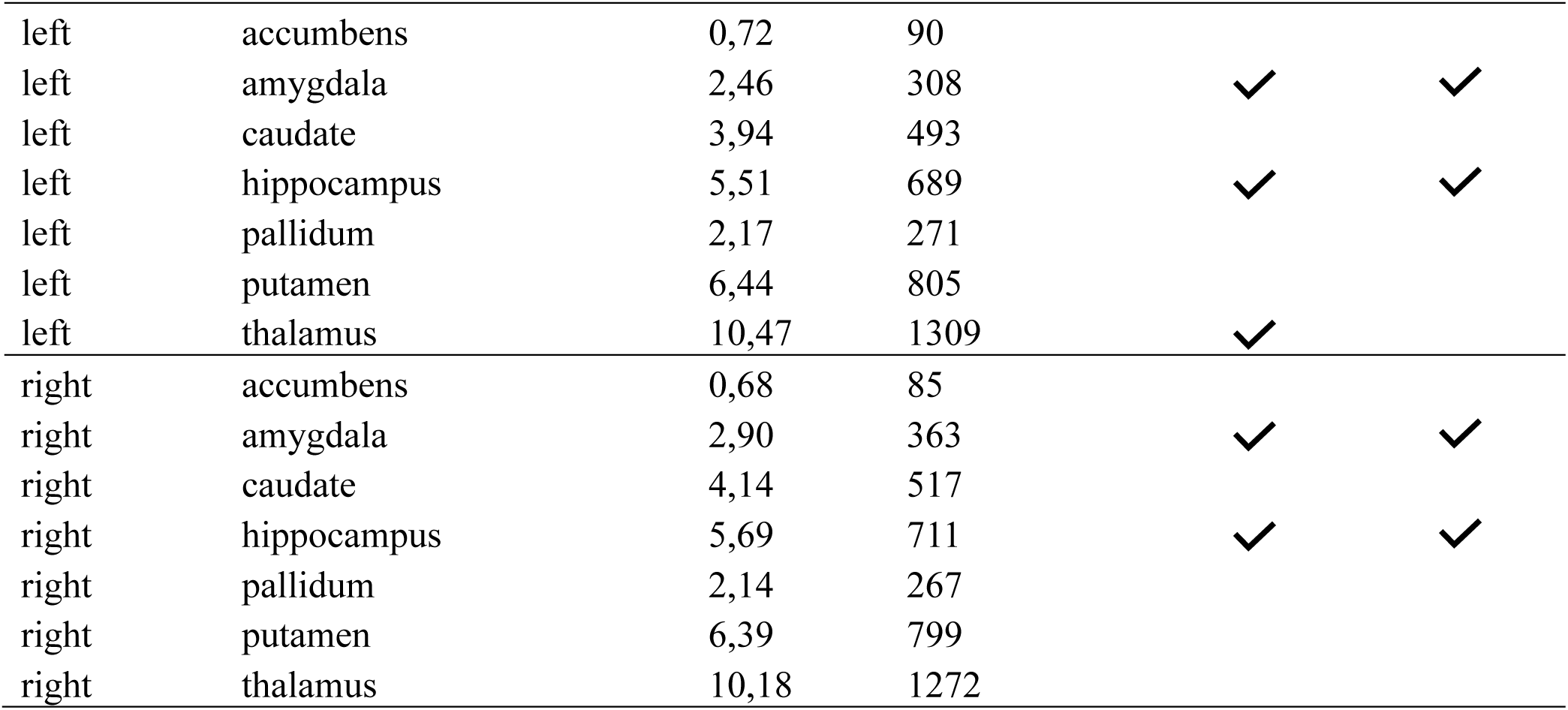
Language regions. The 27 parcels that emerged in the whole-brain GSS analysis (<u>Methods</u>) and the 14 subcortical parcels from Harvard-Oxford Subcortical atlas (Desikan et al., 2006). The parcel sizes are reported in cm^3^ (referring to volumes in the *analysis space*, see <u>Methods</u>) and the number of voxels (2×2x2 mm resolution); individual fROIs are 10% of the parcel size (<u>Methods</u>). The remaining two columns mark the parcels that contain language-responsive and language-selective fROIs (see <u>Results</u> for definitions).

Because the BOLD signal is weaker in brain areas that are further away from the receiver coil, we complemented our functional parcels with a set of 7 bilateral subcortical parcels from the Harvard-Oxford Subcortical atlas (Desikan et al., 2006), to the exclusion of the brain stem, ventricles, and white matter.

### Estimation of response to the three tasks in the individual language fROIs

Within each parcel, we defined a subject-specific functional ROI (fROI) by selecting the 10% of voxels showing the strongest response to the language localizer contrast (*Sentences > Nonwords*) (see **Figures 2** and **3** for sample fROIs; see **Figure OSF4** for evidence that the results are near-identical if the fROIs are defined as voxels that pass a fixed whole-brain significance threshold of p<.001). Subsequently, we estimated the responses in these fROIs to the conditions of the auditory language localizer (*Intact speech* and *Degraded speech*) and the spatial working memory task (*Hard spatial WM* and *Easy spatial WM*). To estimate the responses to the *Sentences* and *Nonwords* conditions, we relied on an across-runs cross-validation procedure, which ensures that different subsets of the data are used for fROI definition and response estimation (e.g., Kriegeskorte et al., 2010).

### Statistical analyses

The data were analyzed using linear mixed-effect models as implemented in the lmerTest package in R (Kuznetsova et al., 2017). We ran a separate analysis for each fROI comparing the critical and baseline conditions in each task: the reading-based language localizer, the auditory language localizer, and the spatial working memory task using the following formula:

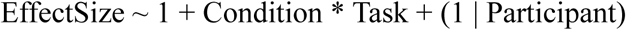

The categorical predictor of *Condition* was deviation-coded using the treatment contrast which specified the control condition of each task as the baseline (the reading-based language localizer: *Sentences* = 1, *Nonwords* = 0; the auditory language localizer: *Intact speech* = 1, *Degraded speech* = 0; the spatial WM task: *Hard WM* = 1, *Easy WM* = 0). The categorical predictor of *Task* was deviation-coded using the treatment contrast and specifying the reading language localizer as the baseline.

Subsequently, we ran three post-hoc pairwise comparisons to test (i) differences between the critical and control conditions for each task (i.e., *Sentences* > *Nonwords*, *Intact speech* > *Degraded speech*, and *Hard WM* > *Easy WM*); (ii) whether in each task the responses to the critical and control conditions significantly differed from 0 (e.g., *Sentences > 0* and *Nonwords > 0* in the reading language task); and (iii) whether the responses to the critical condition significantly differed between a) each of the two language conditions (i.e., *Sentences* > *Intact speech*) and b) the non-linguistic Hard WM condition (i.e., *Sentences > Hard WM*, and *Intact speech > Hard WM*). We do not apply a correction for multiple comparisons (regions) here because we are asking an independent question for each region (rather than asking whether the effect holds in *any* of the regions). Besides, the approach already has internal replicability “checks” built into it, including an across-runs cross-validation procedure for the reading-based contrast and generalization to the auditory language contrast. That said, most of the critical effects would survive a correction for the number of regions given the size of our sample (all GSS and subcortical regions and majority of regions evaluated within standard atlases).

It is important to note that although different subsets of participants completed the three tasks (all n=772 completed the reading-based language localizer; n=489 of the 772 additionally completed the spatial WM task; and n=86 of the 489 additionally completed the auditory language localizer; **Figure 1A**), linear mixed-effects models are well-suited to handle such data imbalances (Little & Rubin, 2019). In particular, this approach efficiently estimates the effects of interest while controlling for data sparsity (assuming greater uncertainty where data are sparse by implementing partial pooling strategy, which “shrinks” the intercepts of participants with less data points towards the population mean more than those of participants who completed all tasks) and is robust under the assumption that the missing data are missing at random, as is reasonable here. In addition, this approach has the advantage of using all available data compared to analyzing each task separately (Snijders & Bosker, 2011; Brown, 2021). Nevertheless, to establish the statistical robustness of our results, we additionally report the results for the subset of the n=86 participants who completed all three tasks and thus where the effects could be estimated with a full within-participant design (**Figure OSF5**).

Finally, although the three functional sequences used in this dataset (see <u>fMRI data acquisition</u>) differ in some acquisition parameters which may be reflected in BOLD responses, we did not explicitly model sequence differences given that (1) all contrasts of interest are within-task, within-run comparisons between conditions, and all effect size estimates come from fROIs that are defined using functional data acquired with the same sequence as the data used for BOLD response estimation. Therefore, any sequence-specific biases signal acquisition would affect all conditions equally and thus cancel out; and (2) the mixed models include by-participant random intercepts, which should absorb any residual between-participant variability in overall signal intensity or SNR.

### Estimation of the language system size

To estimate the size of the language system, we first used individual-level *t*-maps to extract the number of voxels that pass a significance threshold of p<.001, uncorrected, for *Sentences > Nonwords* contrast within the union of all core and extended parcels. To convert this value into cubic centimeters, we multiplied the number of voxels by their size (i.e., 8mm^3^, given that all functional data were resampled to the 2×2x2mm resolution). It is important to note that MNI template tends to be larger than a typical brain. For example, in our data the amount of grey matter in the MNI-normalized probability map was estimated to be, on average, 942cm^3^ (SD = 76cm^3^, min = 548cm^3^, max = 1112cm^3^), whereas in the native space, it was, on average, 748cm^3^ (SD = 74cm^3^, min = 483cm^3^, max = 998cm^3^). As a result, the absolute volume estimates (in cm^3^) in the analysis space that we report may be overestimates of the native-space tissue volumes.

To estimate the proportion that the language system, including the cortical and cerebellar components but excluding the subcortical ones, occupies relative to all grey matter (cortical, cerebellar, subcortical, and brain stem), we used the anatomical segmentation images produced by SPM during preprocessing. In particular, we used the individual grey matter probability maps normalized to the MNI space (wc1* images). Although individual maps that are additionally modulated with a Jacobian determinant (mwc1* images) account for normalization-induced distortions and are generally preferable for estimation of grey matter volume in anatomy-focused analyses (such as voxel-based morphometry), we used non-modulated maps as we wanted to ensure consistency with the analysis space of our functional data. For each participant, we estimated the grey matter amount as the sum of voxelwise grey matter probabilities multiplied by voxel volume.

### Comparison with Standard Cortical Atlases

To facilitate comparisons with other studies, where standard anatomical atlases are often used, we evaluated language responses in three cortical atlases: 1) the Desikan-Killiany-Tourville atlas (Klein & Tourville, 2012), 2) the Harvard-Oxford Cortical atlas (Desikan et al., 2006), and 3) the multi-modal Glasser atlas (Glasser et al., 2016) (see **Figure S5A** for information on the overlap between the functional and anatomical cortical parcels). The aim of these analyses is to evaluate the usefulness of standard atlases as group-constraints for subject-specific fROIs and not making any claims about the relationship of the functional language areas and macroanatomical landmarks.

To create a volume-based DKT parcellation, we ran the cortical reconstruction (using the recon-all Freesurfer standard pipeline) on the MNI template (IXI549Space), which was used in this study to normalize each participant’s data. The resulting parcels (the aparc.DKTatlas+aseg.mgz file) were converted to the NIFTI format and smoothed with a 2 mm FWHM Gaussian kernel. The smoothing step is necessary to slightly increase the spatial extent of the parcels (because they are aligned to the cortical surface, they are relatively thin) as well as accommodate potential small differences between the surface space and the volumetric space. The Harvard-Oxford atlas is volume-based, so no transformation was needed. Finally, for the Glasser atlas, the MNI-based volumetric version was downloaded from https://figshare.com/articles/dataset/HCP-MMP1_0_projected_on_MNI2009a_GM_volumetric_in_NIfTI_format/3501911 and resliced to the 2mm IXI549Space in SPM.

For each atlas, we performed two analyses: an analysis where the whole parcel is used as the ROI (an identical set of voxels is used across participants), and an analysis where within each parcel, we define a subject-specific fROI, as we did in the main analysis (by selecting the 10% of voxels showing the strongest response to the language localizer contrast). Statistical analyses were performed on the responses extracted from these two kinds of ROIs (anatomical group-level ROIs and individual fROIs) using the same approach of linear mixed-effects models, as described above.

### Code accessibility

All data, including 3D brain contrast maps (both smoothed and unsmoothed), parcels, and data produced by the functional localization analysis toolbox (spm_ss), as well as the code necessary to reproduce all analyses and plots presented in this paper are available at: https://osf.io/7594t/.

## Results

We considered a region to contain a fROI **responsive to language** if the following conditions were met: 1) the response to both of the language contrasts (*Sentences* > *Nonwords* and *Intact* > *Degraded speech*) is positive and significant (p<.05), and 2) the response to both of the critical conditions (i.e., *Sentences* for the reading language localizer and *Intact speech* for the auditory language localizer) falls significantly above the fixation baseline. We further considered a region to contain a fROI **selective for language** if, in addition to the abovementioned conditions, the critical language conditions (*Sentences* or *Intact speech*) elicited a reliably greater response than the critical condition for the non-linguistic task (i.e., *Hard WM*). Note that here, we only evaluate selectivity for language relative to one task: a non-linguistic working memory task. This approach allows us to determine whether a brain region responds to any cognitive task, or whether the response reflects processing required for language comprehension (through reading and listening) but not for performing the working memory task. This limited definition of selectivity already rules out some of the language-responsive regions as supporting computations that are not specific to language (see <u>Results</u>), but it is of course possible that some of the regions found here to be selective will later be found to not be selective relative to other non-linguistic tasks (such as processing music; or processing non-verbal communicative signals; or processing non-linguistic meanings).

### The extended set of language-selective regions

Twenty-two of the 27 parcels identified by the whole-brain GSS analysis (<u>Methods</u>) contained fROIs that responded to language across modalities, with the exception of the three parcels in the occipital cortex, the parcel in the left precuneus / posterior ACC, and the parcel in the left parahippocampal gyrus (the fROIs in these parcels either did not show a significant response to one of the language contrasts or responded at or below baseline to one or both of the critical language conditions). Of the 22 language-responsive regions, 20 (including 5 core LH regions and their RH homotopic regions) showed selective responses to language with little or no response to the demanding non-linguistic task; both of the exceptions were located in the cerebellum (**Figures 2** and **3**, **Table 1**; see **Table S1** for the details of the statistical tests; for completeness, see **Figure S1** for the profiles of the regions excluded due to small parcel size). Note that in addition to these language-selective areas, most of these parcels contain areas that respond strongly to the non-linguistic task and not to the language tasks (see **Figure S3**), in line with past evidence that distinct functional areas often lie near each other within the same anatomical structure (e.g., Fedorenko et al., 2012; Braga et al., 2020; Du et al., 2024)—a finding that reinforces the importance of functionally defining the relevant region within each individual.

In addition to the core left lateral frontal and lateral temporal regions and their homotopic regions—which replicate prior findings (Mahowald & Fedorenko, 2016; see Fedorenko et al., 2024 for a review) and which we report here for completeness—the extended language-selective network (**Figure 3**) included regions in the bilateral temporal pole, in the left ‘basal temporal language area’ (BTLA; Lüders et al., 1986; see Li et al., 2024 and Salvo et al., 2025 for concordant evidence using a similar functional localization approach to the one used here), in the left precuneus, in three regions in the medial frontal cortex (spanning different sections of the SFG and the frontal pole), and in three cerebellar regions (left Crus II / VIIb, right Crus I / II / VIIb, and right VIIb / VIIIa; see Casto et al., 2025a for concordant evidence).

It is important to note that even with these additional regions, language-selective cortex takes up a relatively small amount of grey matter volume. Although the parcels are, by design, large (to capture inter-individual variability in the precise locations of functional areas), the individual fROIs constitute only a small fraction of each parcel. For example, voxels that show a significant (p<.001) *Sentences>Nonwords* effect across all our 20 parcels (10 core and 10 non-canonical) add up to 22.1 cm^3^, on average (SD=12.8 cm^3^), which corresponds to only 9.4% of the volume of the group language parcels, and constitutes ∼3.5% of the grey matter volume. Note that this estimate is based on non-modulated grey matter maps—to match the analysis space of the functional data (see <u>Methods</u>)— and thus may slightly over- or under-estimate true anatomical volumes, but it should still fall within the right ballpark showing that the language system takes up only a small fraction of grey matter. The volume estimate is also similar when taking top 10% of language-responsive voxels in each parcel, and language selectivity declines steeply beyond this threshold (see **OSF6**). An average of 15.2 cm^3^ (SD=8.2 cm^3^) is taken up by the five core left-hemisphere fROIs (defined within the following parcels: AntTemp, PostTemp, IFGorb, IFG, and MFG), and 3.6 cm^3^ (SD=3.7 cm^3^)—by their right-hemisphere homotopes.

In addition to the cortical language areas, 4 of the 14 parcels that we examined in the Harvard-Oxford Subcortical atlas contained language-selective fROIs: bilateral hippocampi and bilateral amygdalae (**Figure 4**, **Table 1**). One additional parcel (the left thalamus) contained a language-responsive, but non-selective fROI (see **Table S1** for the details of the statistical tests).

**Figure 4.**
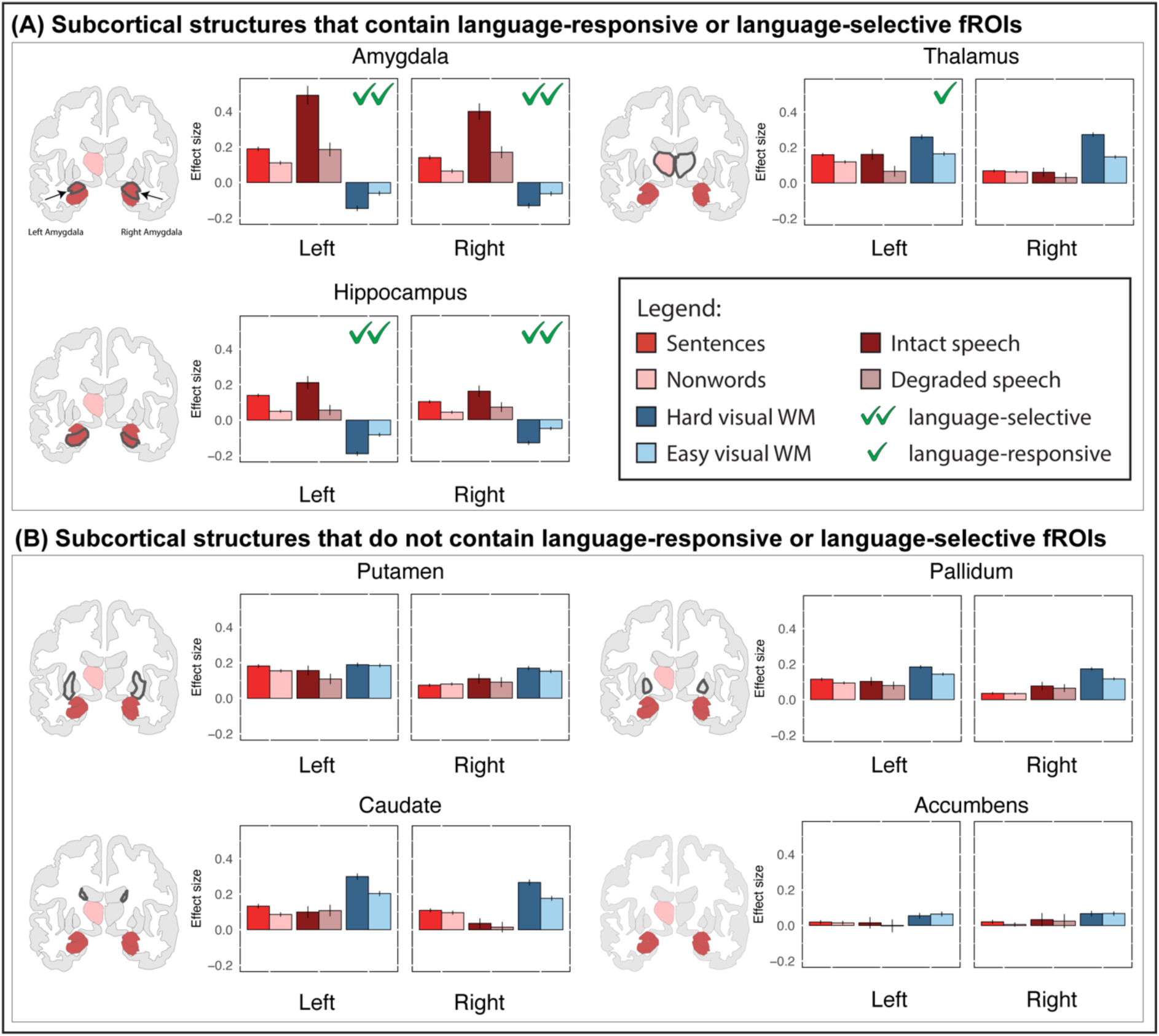
Language responsive regions within the parcels in the Harvard-Oxford Subcortical atlas. For each region (defined based on individual activation maps; <u>Methods</u>), we show responses to the reading-based and auditory language localizers (red bars; darker bars correspond to the critical conditions) and the non-linguistic spatial working memory task (blue bars; the darker bar corresponds to the harder condition). For the reading-based localizer, the responses were estimated using independent runs of the data. A) Parcels that contain language-responsive fROIs have a green checkmark; the ones where the fROIs are additionally language-selective have two green checkmarks. B) The parcels that do not contain language-responsive fROIs.

The full set of 27 cortical parcels, in both the MNI and the fsaverage spaces, is available on OSF (https://osf.io/7594t/; note that the fsaverage version was created by converting the volumetric set to the fsaverage space, not based on statistical maps generated in a surface space). Additionally, for the 20 parcels containing language-selective fROIs, we created a left-right symmetrical version, where the left-hemisphere cortical parcels were mirrored onto the right hemisphere (the medial parcels were first split along the brain midline (x = 45)), and the right-hemisphere cerebellar parcels were mirrored onto the left hemisphere. This symmetrical set (also available on OSF at https://osf.io/7594t/ and at https://evlab.mit.edu/parcels) may be useful for defining the language fROIs in atypically lateralized (right-lateralized or bilateral) participants, or for directly comparing the profiles of the left- and right-hemisphere language regions. We also provide a left-right symmetrical version that uses—for the core areas—the parcels derived from a smaller set of participants and used in much past work (e.g., Diachek, Blank, Siegelman et al., 2020; Ivanova et al., 2020; Shain et al., 2020; Schrimpf et al., 2021; Malik-Moraleda, Ayyash, et al., 2022).

### Language responses in three standard brain atlases

To facilitate comparisons with other work, we comprehensively evaluated language responses in three cortical atlases that vary in the granularity of their parcellation. These analyses serve two goals. First, they help identify anatomical areas within which reliable and selective responses to language can be observed within individuals. And second, they highlight the critical importance of individual-level functional localization.

When using the functional localization approach (i.e., selecting—for each individual separately—the top 10% of most active voxels based on their language localizer map), a number of areas emerge as containing language-selective fROIs (**Figure 5**). Full functional profiles of the atlas parcels are presented on **Figure S6** (see **Table OSF2** for details of statistical tests in all regions in the three atlases). In the DKT atlas, 20 of the 72 parcels contain language-selective fROIs: 13 in the left hemisphere and 7 in the right hemisphere. In the Harvard-Oxford Cortical atlas, 42 of the 96 parcels contain language-selective fROIs: 26 in the left hemisphere and 16 in the right hemisphere. Finally, in the Glasser multimodal atlas, 95 of the 380 parcels contain language-selective fROIs: 64 in the left hemisphere and 31 in the right hemisphere (see **Table 2** for the full list of language-responsive and language-selective regions). Across all three atlases, the topography of the parcels containing language-selective fROIs resembles the topography of the functional parcels from the GSS (see **Figure S5A** for information on the overlap between standard atlas parcels and functional language parcels), spanning bilateral frontal and temporal areas, as well as pre-central frontal and inferior parietal areas in the left hemisphere. However, in the most granular, Glasser atlas, 9 of the parcels containing language-selective fROIs showed no or minimal spatial overlap with any of the GSS parcels (see **Figure S2** for the functional profiles of these regions).

**Figure 5.**
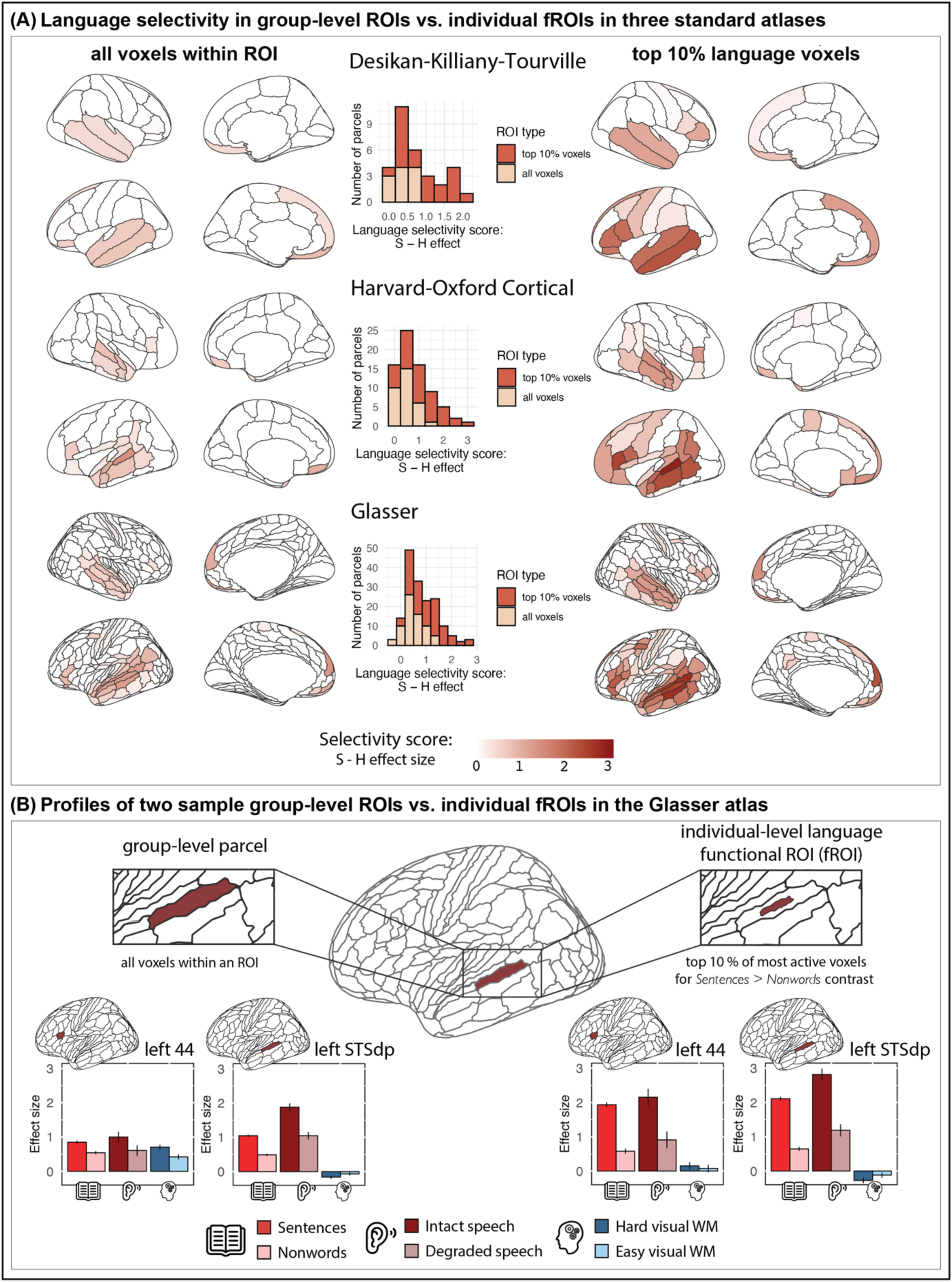
Consequences of using entire group-level anatomical ROIs vs. individually-defined functional ROIs for evaluating language selectivity. A) Across three standard cortical parcellations (rows; top: DKT: Klein & Tourville, 2012; middle: HOC: Desikan et al., 2006; bottom: Glasser: Glasser et al., 2016), we show the selectivity scores (difference between the response to the *Sentences* condition and the *Hard spatial WM* condition) for all regions. Higher selectivity scores are shown with darker red hues (see bottom of the panel for the legend). In the left column, the selectivity scores are based on using group-level anatomical ROIs (all voxels in a parcel are used); in the right column, the selectivity scores are based on using individually-defined functional ROIs (top 10% of voxels are selected based on individual activation maps; the responses to the Sentence condition are estimated in a left-out run of the data; <u>Methods</u>). For each atlas and each ROI definition (group-level vs. individual-level), four views of the brain are shown: right lateral and medial and left lateral and medial. (For a figure showing, for each atlas, effect sizes for the three individual contrasts—*Sentences>Nonwords*, *Intact>Degraded speech*, and *Hard>Easy spatial WM*—see **Figure S6**). In the middle column, we show—for each atlas—a quantitative comparison of the distributions of selectivity scores from the group ROIs vs. individual fROIs. B) For two sample parcels in the Glasser atlas (Glasser et al., 2016)—left 44 and left STSdp (selected based on high overlap with the probabilistic atlas of the language network; Lipkin et al., 2022)—we show the response profiles for the group-level anatomical ROIs (left) and individual-level fROIs (right).

**Table 2.** List of parcels that contain language-responsive and language-selective fROIs in the three standard cortical atlases: the DKT, Harvard-Oxford Cortical, and Glasser atlases. In the Glasser atlas, parcels marked with a degree sign (°) lie at the intersection of two anatomical lobes and parcels marked with a cross lie at the intersection of three anatomical lobes and are thus listed in more than one lobe.

| Parcels in the DKT atlas that contain language-selective* fROIs |  |  |
| --- | --- | --- |
| Lobe | Left hemisphere | Right hemisphere |
| Frontal | Caudal Middle Frontal, Medial Orbitofrontal, Pars Opercularis, Pars Orbitalis, Pars Triangularis, Precentral, Superior Frontal, Lateral Orbitofrontal | Medial Orbitofrontal, Pars Opercularis, Pars Triangularis, Precentral, Superior Frontal |
| Parietal | Supramarginal, Postcentral |  |
| Temporal | Inferior Temporal, Middle Temporal, Superior Temporal | Middle Temporal, Superior Temporal |

| Parcels in the Harvard-Oxford Cortical atlas that contain language-selective fROIs |  |  |
| --- | --- | --- |
| Lobe | Left hemisphere | Right hemisphere |
| Frontal | Frontal Pole, Sup. Frontal Gyrus, Mid. Frontal Gyrus, Inf. Frontal Gyrus Pars Triangularis, Inf. Frontal Gyrus Pars Opercularis, Precentral Gyrus, Frontal Medial Cortex, Juxtapositional Lobule Cortex, Frontal Orbital Cortex, Frontal Operculum Cortex | Inf. Frontal Gyrus Pars Triangularis, Inf. Frontal Gyrus Pars Opercularis, Frontal Medial Cortex, Frontal Orbital Cortex, Juxtapositional Lobule Cortex |
| Parietal | Supramarginal Gyrus Post., Angular Gyrus, Central Opercular Cortex, Parietal Operculum Cortex | Supramarginal Gyrus Post., Angular Gyrus |
| Temporal | Temporal Pole, Sup. Temporal Gyrus Ant., Sup. Temporal Gyrus Post., Mid. Temporal Gyrus Ant., Mid. Temporal Gyrus Post., Mid. Temporal Gyrus Temporooccipital, Inf. Temporal Gyrus Ant., Inf. Temporal Gyrus Post., Temporal Fusiform Cortex Ant., Temporal Fusiform Cortex Post., Planum Polare, Planum Temporale | Temporal Pole, Sup. Temporal Gyrus Ant., Sup. Temporal Gyrus Post., Mid. Temporal Gyrus Ant., Mid. Temporal Gyrus Post., Mid. Temporal Gyrus Temporooccipital, Inf. Temporal Gyrus Ant., Planum Temporale, Temporal Fusiform Cortex Ant. |
|  | **Additional parcels that contain language-responsive (but not language-selective) fROIs** |  |
| Frontal |  | Precentral Gyrus |
| Parietal | Supramarginal Gyrus Ant. |  |
| Temporal | Inf. Temporal Gyrus Temporooccipital |  |
| Occipital | Lat. Occipital Cortex Inf. |  |

| Parcels in the Glasser atlas that contain language-selective fROIs |  |  |
| --- | --- | --- |
| Lobe | Left hemisphere | Right hemisphere |
| Frontal | FEF, PEF, 4, 55B, SFL, 8BM, 10R, 47M, 9M, 9P, 44, 45, 47L, 6R, 8AV, 8BL, IFJA, IFSP, IFSA, 9A, 10V, 10d, 47S, FOP5, S32°, FOP4°, OP4°, OP2-3<sup>†</sup>, FOP1 | 55B, SFL, 47M, 9M, 8BL, 44, 45, IFJA, IFSP, IFSA, 10V, OFC |
| Parietal | 3b, PFCM, PF, OP4°, 31PD°, PGI†, PSL°, OP2-3† | 3B |
| Occipital | PGI†, PHT°, TPOJ3 | PHT°, TOPJ3 |
| Temporal | FOP4°, PEEC°, PGI†, PSL°, STV, RI, TA2, STGA, PBELT, A5, STSDA, STSDP, STSVP, TGD, TE1A, TE1P, TF, TE2P, TPOJ1, TPOJ2, TGV, A4, STSVA, TE1M, PI, PIR°, PHT°, OP2-3† | STV, TA2, STGA, A5, STSDA, STSDP, STSVP, TGD, TE1A, TE1P, TPOJ1, TPOJ2, TGV, A4, STSVA, TE1M, PHT° |
| Limbic** | S32°, 31PV, PEEC°, 31PD°, PIR° H, pOFC |  |
| **Additional parcels that contain language-responsive (but not language-selective) fROIs** |  |  |
| Frontal | SCEF, 6D, 6V, IFJP, P9-46V, I6-8, 8C | 6r, 6V, FEF, IFJP, PEF, SCEF |
| Parietal | 1 | 1 |
| Temporal | PHA3 | TF |
| Occipital | MST, FST |  |

\* All DKT language-responsive regions are also language-selective.
\*\* Lobe segmentation in the Glasser parcellation follows PALS\_B12 atlas (Van Essen, 2005;
https://surfer.nmr.mgh.harvard.edu/fswiki/PALS_B12) template (using FreeSurfer);
° denotes regions straddling two lobes;
† denotes regions straddling three lobes.

In stark contrast, when using the whole parcels as ROIs (taking all voxels within an ROI, with no participant-specific functional masking), many fewer areas emerge as language-selective, and in the ones that do, the degree of selectivity is substantially lower compared to the corresponding fROIs (see the histograms in **Figure 5B** for a quantitative comparison). For example, in the DKT atlas, which is the least granular (with ROIs corresponding to entire cortical gyri in many cases), only 11 regions satisfy the selectivity criterion when the entire regions are used as ROIs (cf. 20 regions when using individual-level fROIs). The selectivity of these regions (the difference between the response to *Sentences* vs. to the *Hard spatial WM* condition) is low (2-3 times lower than when using fROIs). Further, some regions, especially in the frontal cortex, show mixed selectivity (responses to both the language contrasts and the spatial WM contrast), when in reality, little or no overlap exists between these contrasts within any given individual (**Figure 5B**).

The picture is similar for the other two atlases: fewer regions emerge as language-selective, and in the ones that do, selectivity is severely underestimated. For example, in the Harvard-Oxford atlas, which has an intermediate level of granularity, in the two most selective group-level ROIs, the difference between the *Sentences* condition and the *Hard spatial WM* condition is 1.39 (left posterior superior temporal gyrus) and 1.03 (left anterior superior temporal gyrus); the corresponding effect sizes in the individually-defined fROIs within these regions are 2.80 and 1.97. Similarly, even in the most granular atlas (Glasser et al., 2016), the selectivity of the individual fROIs is substantially higher (average across all selective fROIs: 1.02, SD = 0.65, range 0.05-2.67; compared to the average of the group-level ROIs (average: 0.50, SD = 0.39, range = −0.37 – 1.33; **Figure 5A**).

These differences in selectivity can be straightforwardly explained by i) inter-individual variability in the precise locations of language fROIs within the larger, anatomical regions (see **Figure S5B** for examples), and ii) the presence of domain-general areas that respond robustly to demanding cognitive tasks in the vicinity of the language areas (**Figures S3** and **S4**). In particular, a set of voxels that is the same across participants in a common template space is bound to include—for any given individual—some non-localizer-responsive voxels, leading to a lower response to the *Sentences* condition, and it may further include voxels that belong to the nearby Multiple Demand network (Fedorenko & Blank, 2020; Braga et al., 2020), leading to a higher response to the working memory task conditions (see e.g., Nieto-Castañón & Fedorenko, 2012 for discussion).

## Discussion

Many brain areas outside the left fronto-temporal network have been previously implicated in language processing (**Table OSF1**). Here, we systematically searched for language-responsive areas across the brain. To do so, we a) used a paradigm that isolates language processing from perceptual, motor, and task-related cognitive demands (Fedorenko et al., 2010), b) included a large dataset (n=772 participants), c) examined language responses across modalities, and d) evaluated language selectivity relative to a demanding cognitive task. In addition to the core left hemisphere areas and their homotopes, we identified 17 areas (7 cortical and 10 subcortical, including 5 cerebellar) that respond to auditory and written language, with 14 of these areas showing selectivity for language relative to a demanding non-linguistic task.

### The contributions of non-canonical language areas to language and cognition

Establishing that a brain area responds to language across modalities and not during a non-linguistic task ensures that its activity is not driven by perceptual or general cognitive demands. Still, the precise contributions of these language-responsive and language-selective areas remain to be discovered. Past studies may hold some clues, but relating our results to prior work is challenging: we cannot be certain that the areas we found here correspond to those identified in prior work, because macroanatomy is a poor guide to function in the association cortex (Frost & Goebel, 2012; Tahmasebi et al., 2012; also, **Figure S5B**). Adopting the functional localization approach has led to several robust and replicable findings about the core language network, including its selectivity for language and its role in linguistic computations (see Fedorenko et al., 2024 for a review). The extended language areas we identified here constitute new targets for similar systematic investigations, which should i) evaluate the selectivity of these areas relative to other functions that bear similarities to language, ii) probe sensitivity of these areas to linguistic demands associated with lexical and combinatorial processing (in controlled and naturalistic paradigms), and iii) directly compare these areas’ contributions to those of the core language areas.

The non-selective language-responsive areas may still meaningfully contribute to language processing. We have identified three such areas: two in the right cerebellum (Casto et al., 2025a) and one in the left thalamus. Higher spatial resolution approaches may eventually reveal that distinct neural sub-populations create the appearance of mixed selectivity. However, if the mixed selectivity is real, it may suggest that these regions play an integratory role, perhaps combining inputs from the core language areas and other networks. Indeed, both the cerebellum and the thalamus have been hypothesized to serve as information integrators (Barbas et al., 2013; Theofanopoulou & Boeckx, 2016; Wolpert et al., 1998).

### Activation during a “language task” may not always reflect linguistic processing

Some past studies have reported neural activity during language tasks in brain areas that do not show language-responsiveness in our study. However, activation during “language tasks” (i.e., tasks that use verbal materials) may not always reflect linguistic processing. Many language paradigms tax both linguistic and general cognitive demands, so a response in such paradigms could reflect the latter. For example, Thibault et al. (2021, 2025) have argued for overlap between syntactic processing and tool use in the basal ganglia. However, both the syntactic and tool-use tasks in those studies require cognitive processes beyond syntactic and tool-use-related computations. We show that the language paradigms that are not confounded with task demands do not consistently engage any basal ganglia structures (**Figure 4B**): whereas the reading language contrast elicits a small response in the left caudate and putamen, these responses do not generalize to the auditory language contrast, which rules out the possibility that these brain areas are engaged in language processing. These findings suggest that the syntactic complexity manipulation in Thibault et al.’s studies may be engaging these areas due to general cognitive demands and not by virtue of syntactic computations (see **Figure S4** for evidence that basal ganglia areas respond strongly to task demands).

Similarly, Seydell-Greenwald et al. (2023) report responses to auditory language tasks in the primary visual cortex (V1) of sighted individuals. We do not replicate this finding:, the weak response in occipital areas to the reading-based contrast (mostly due to greater deactivation during nonword reading) does not generalize to the auditory modality, which rules out the possibility that V1 implements linguistic computations (cf. evidence of V1 supporting language processing in blind individuals; Bedny et al., 2011; Lane et al., 2015; Pant et al., 2020; Czarnecka et al., 2025). The effects observed in Seydell-Greenwald et al. (2023) might be explained by the fact that their contrasts are confounded with general cognitive demands: one contrast entails listening to sentences and answering questions about them vs. listening to reversed speech and responding to a beep (see Ozernov-Palchik, O’Brien et al., in press for additional discussion of difficulty confounds in this paradigm), and the other contrast compares listening to words and pressing a button for catch trials (thus requiring attention to the task beyond the lexical processing demands) vs. a low-level baseline. Attentional modulation of V1 is well-established (Somers et al., 1999), so the observed responses plausibly reflect attention-related engagement. To argue that a brain area supports linguistic computations, one needs to ensure that it responds to language across tasks and modalities but also that the language task is not confounded by difficulty or attentional demands. Previous work has established that the core language areas are relatively insensitive to task demands, showing similarly strong responses during passive comprehension and paradigms that include tasks (Fedorenko et al., 2010; Diachek, Blank, Siegelman et al., 2020; Gao et al., 2025). Here, we extend these findings to several non-canonical language areas but also question some past findings of “language” responses that have alternative explanations.

### A small and well-defined subset of the brain implements language processing

Despite ample evidence for functional specialization in the brains of humans (Kanwisher, 2010) and non-human animals (Tsao et al., 2006), some continue to argue against the idea of stable structure in the brain, emphasizing the distributed, dynamic, and interactive nature of cognitive processes, including lnguage (e.g., Pessoa, 2022; Forkel & Hagoort, 2024; Drijvers et al., 2025). Deep engagement with this debate is beyond the scope of this article, but two points are worth clarifying. First, the fact that many areas—sometimes in distant parts of the brain—are engaged by language comprehension does not imply that the “entire brain” supports this function. Although the extended language network spans almost every major component of the brain, within each component, language regions occupy a small fraction of brain tissue. Second, linguistic inputs can unquestionably engage many brain regions beyond those that specifically support language processing: vivid descriptions of faces or scenes can engage category-selective visual areas, a story about a misunderstanding can engage the Theory of Mind network, and a horror story can engage the amygdala (see Casto, et al., 2025b for discussion). However, all these brain regions can also be engaged by non-linguistic inputs. The ability of a brain region to be engaged by language does not make it a “language region” any more than its ability to be engaged by visual inputs makes it a “visual region”. Furthermore, the fact that the language network needs to interact with other brain areas does not undermine its functional distinctness from those areas and its special role in language processing. As long as different components within the language network interact more strongly with one another than with other networks—for which ample evidence exists (Blank et al., 2014; Braga et al., 2020; Du et al., 2024, 2025; Shain & Fedorenko, 2025)—the language network and other cognitive networks can be treated as meaningfully distinct objects of study (Simon, 1962).

### Parcels derived from functional data have advantages over anatomical parcels for constraining individual fROIs

For completeness and ease of comparison with past studies, we explored the possibility of using standard anatomical atlases to constrain individual fROIs (rather than the parcels derived from a group-level representation of brain activity; Fedorenko et al., 2010; Julian et al., 2012). Examining individual activation maps (or maps derived from functional connectivity patterns: Braga et al., 2020; Du et al., 2024, 2025; Shain & Fedorenko, 2025) against standardized brain parcellations reveals two issues. First, individual topographies often do not align with the boundaries of the atlas areas: a contiguous functional region may get broken up by a boundary, or—for finer-grained atlases—may get assigned to different atlas areas across individuals because of inter-individual topographic variability (see **Figure S5B** for examples). For studies focusing on a particular functional network, we therefore recommend functional parcels over anatomical/multimodal atlases. Critically, however, whichever approach is chosen, the regions of analysis should be defined functionally within individuals (using localizers or functional-connectivity-based parcellations) to avoid conflating nearby networks at the group level (**Figure 4**; see **Figure S3** for evidence that most language parcels include regions of the Multiple Demand network). And second, using large, anatomical ROIs creates an illusion that language processing engages large chunks of the cortex, while in reality these extensive regions contain (sometimes multiple) small language-responsive regions (see **Figures 2A and 3A** for illustration of the size of individual-level fROIs relative to the parcels and **Figure 5** and **Figure S5A** for comparison of the extent of language responsive areas in coarser- vs. finer-grained parcellations).

In conclusion, we have comprehensively searched for language-selective areas across the brain and identified several new components of the extended language network. This work lays the groundwork for future investigations of these new components, including their similarities to and differences from the core language areas.

## Acknowledgements

We would like to acknowledge the Athinoula A. Martinos Imaging Center at the McGovern Institute for Brain Research at MIT, and its support team (Steve Shannon and Atsushi Takahashi). We would also like to thank Moshe Poliak for statistical consultations as well as former and current EvLab members for contributions to the fMRI data collection efforts over the years. This work was partially supported by a grant from the Simons Foundation to the Simons Center for the Social Brain at MIT. AW was supported by a postdoctoral fellowship from the Simons Center for the Social Brain. CC as supported by the Kempner Institute for the Study of Natural and Artificial Intelligence at Harvard University. EF was additionally supported by research funds from the McGovern Institute for Brain Research, the Department of Brain and Cognitive Sciences, and the MIT Siegel Family Quest for Intelligence.

## Author Contributions

Conceptualization: AgW, EF; Methodology: all authors; Software: AgW, AaW; Validation: AaW; Formal analysis: AgW, AaW; Investigation: all authors; Data curation: all authors; Writing – original draft: AgW, EF; Writing – review & editing: AaW, CC, SH, BL; Visualization: AgW, AaW, SH; Supervision: EF; Project administration: AgW, EF.

