## supplemental materials for "The extended language network: Language-responsive brain areas whose contributions to language remain to be discovered"

### Supplemental Information for Wolna et al. (2025)

Section 1: **Functional profiles of additional language-responsive regions not included in the main analyses.**

***Figure S1.1.*** Functional profiles of the language regions that emerged in the GSS analysis but were excluded due to small size.
***Figure S1.2.*** Functional profiles of the language regions in the Glasser atlas that show minimal overlap with the GSS language parcels.

Section 2: **Domain-general Multiple Demand (MD) regions lie in close proximity to many of the language regions.**

***Figure S2.1.*** Functional profiles of the MD fROIs within the new, extended set of language parcels.
***Figure S2.2.*** Functional profiles of the MD fROIs within the parcels of the Harvard-Oxford Subcortical atlas.

Section 3: **The relationship between language activations and standard anatomical atlases.**

***Figure S3.1.*** The relationship between language activations and standard anatomical atlases.
***Figure S3.2.*** Language responsiveness and selectivity of the language fROIs defined within the parcels in three standard cortical atlases.

***Table S1.*** Statistical results for the key contrasts across the three paradigms (reading-based language localizer, auditory language localizer, and spatial working memory) in the language regions a) from the GSS analysis, and b) from the Harvard-Oxford Subcortical atlas.

For some **additional control analyses**, please see the materials included on OSF (https://osf.io/7594t/):

**OSF 1:** The Sentences > Nonwords contrast subsumes brain areas engaged during nonword processing.
**OSF 2:** The results are robust to preprocessing choices (template vs. native space, and volume vs. surface).
**OSF 3:** The functional topography of the language network is similar across different ways of creating the probabilistic language atlas.
**OSF 4:** The results are robust to the fROI definition procedure.
**OSF 5:** The results are robust to participant selection.

### Section 1: Functional profiles of additional language-responsive regions not included in the main analyses.

In the main text, we report functional profiles for 27 language regions identified with the GSS parcellation. The parcellation, however, yielded 33 regions in total, 6 of which were smaller than 150 voxels. Here, for completeness, in **Figure S1.1.**, we include functional profiles for 5 of these 6 regions; the remaining parcel was only 5 voxels in size and therefore could not be analyzed with our standard approach of defining fROIs as the top 10% of most responsive voxels. We also provide the full set of 33 parcels from the GSS parcellation on OSF (https://osf.io/7594t/files/).

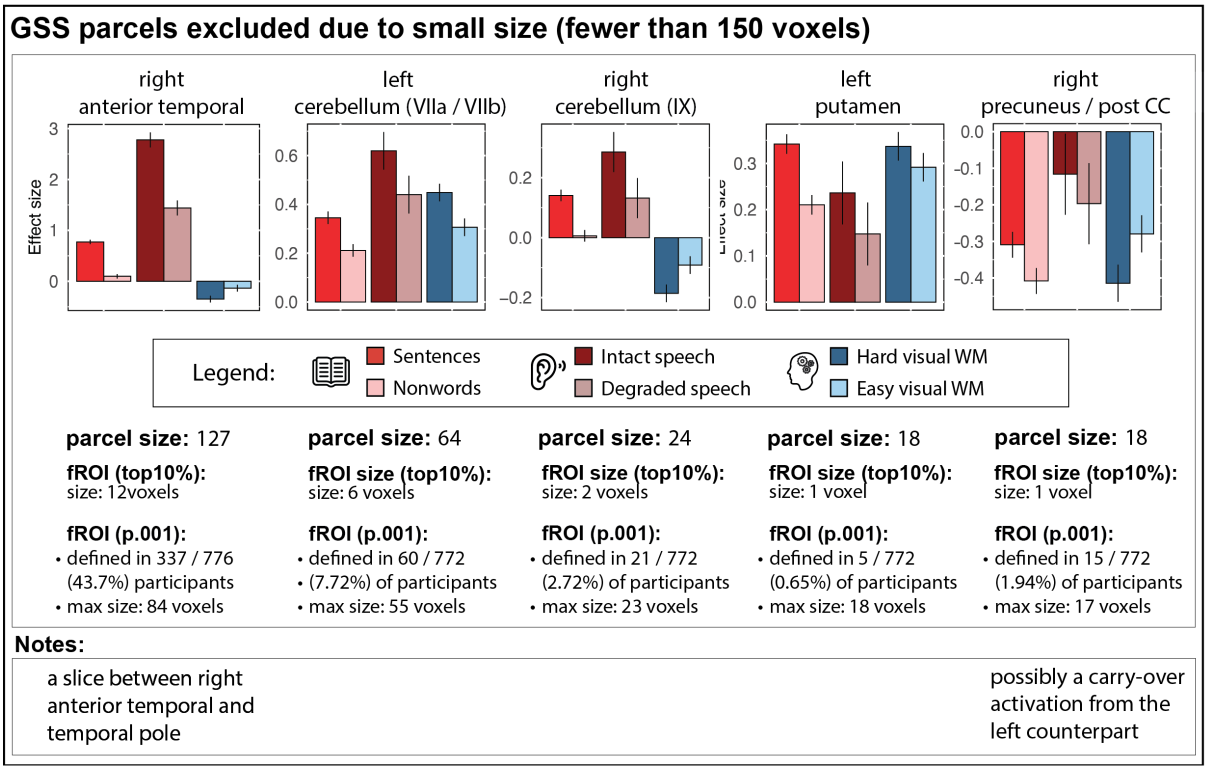


**Figure S1.1. Functional profiles of the language regions that emerged in the GSS analysis but were excluded due to small size.** Five parcels were excluded from the main analysis due being smaller than 150 voxels in size. We show the functional profiles of these regions and provide details on the size of the parcel and the fROI (defined using the top 10% approach, or a fixed threshold of $p < .001$, uncorrected); for the latter, we provide the proportion of participants in whom the fROI could be defined in our sample (i.e., contained at least 1 supra-threshold voxel) and the maximum size. In each bar plot, the bars correspond to the two conditions of the the reading-based and auditory language localizers (red bars; darker bars correspond to the critical conditions) and the two conditions of the spatial working memory task (blue bars; the darker bar corresponds to the harder condition). Note that the y-scale differs across regions.

All but one region (the left putamen) meet our criteria for language responsiveness, and two meet our criteria for language selectivity (the right anterior temporal region and the right cerebellar region). However, all these fROIs contain very few voxels when defined using the top 10% approach and can only be defined in a fraction of participants when using a fixed, p<.001 threshold. As a result, we chose to exclude these regions from the main analyses.

Further, in our analyses of language fROIs defined within the anatomical parcels in standard cortical parcellations (DKT: Klein & Tourville, 2012; HOC: Desikan et al., 2006; Glasser: Glasser et al., 2016), we found 9 parcels in the Glasser atlas that contained language-selective fROIs but showed no or minimal (below 10%) overlap with the GSS language parcels. These fROIs' functional profiles are presented in **Figure S1.2.** All of these parcels are small in size, and the differences between the critical and control conditions in the language paradigms are often small and may not come out as reliable in more typical sample sizes.

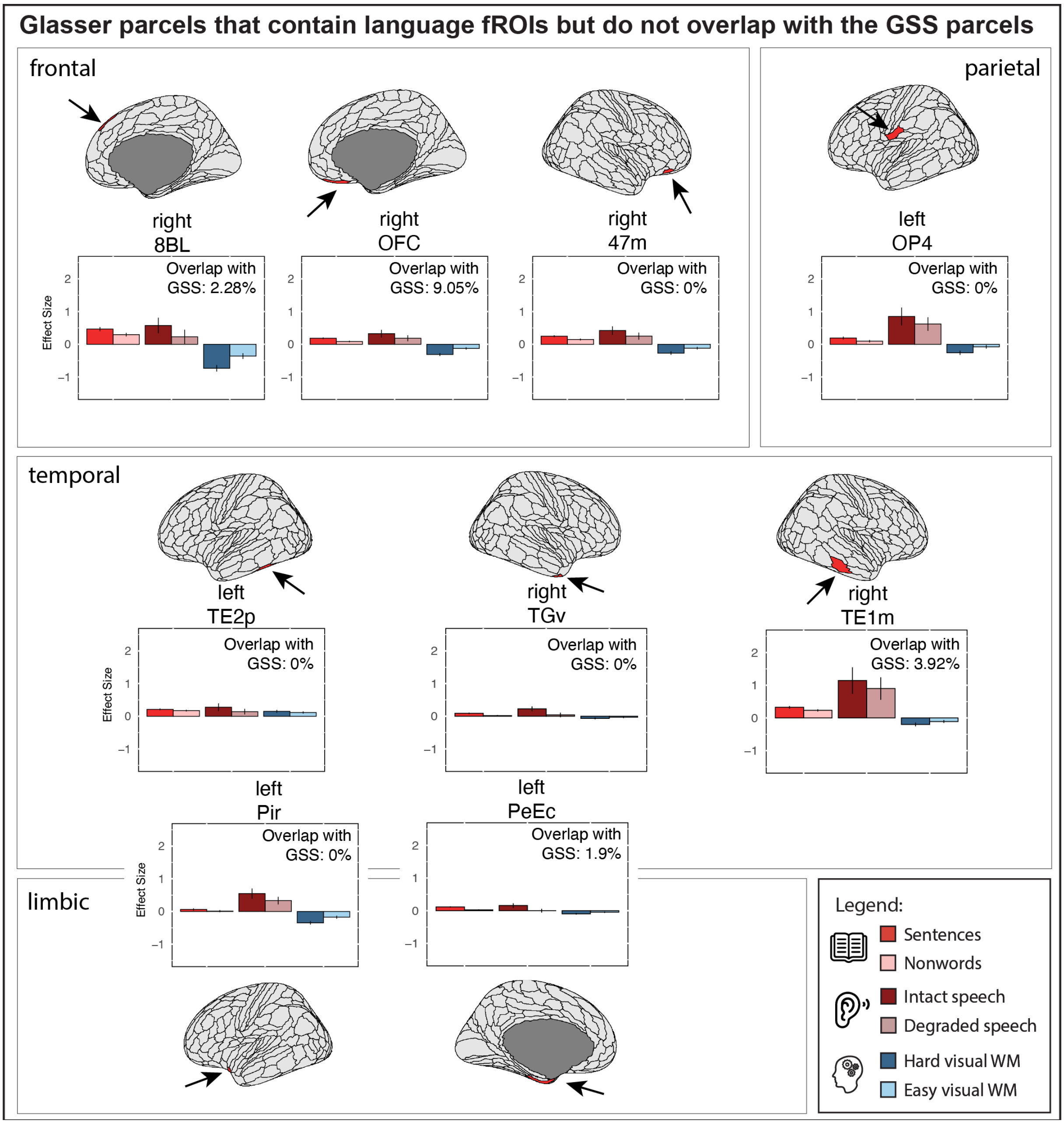


**Figure S1.2. Functional profiles of the language regions defined within the Glasser parcels that show minimal overlap with the GSS language parcels.** We show the functional profiles of these regions (defined using the top 10% approach). In each bar plot, the bars correspond to the two conditions of the the reading-based and auditory language localizers (red bars; darker bars correspond to the critical conditions) and the two conditions of the spatial working memory task (blue bars; the darker bar corresponds to the harder condition). Spatial overlap with the GSS parcels is shown on each plot in the top right corner. Separate panels correspond to brain lobes that the parcels fall into, with plots falling at the intersection of two panels being assigned to two lobes. Lobe segmentation follows PALS_B12 atlas (Van Essen, 2005; https://surfer.nmr.mgh.harvard.edu/fswiki/PALS_B12) template (using FreeSurfer).

#### Section 2: Domain-general Multiple Demand (MD) regions lie in close proximity to many of the language regions.

In the main text, we report analyses of language fROIs (i.e., fROIs defined by the language localizer contrast: *Sentences* > *Nonwords*) in the new set of extended language parcels. Here, we ask whether the language parcels include fROIs that reliably respond to general task demands. To do so, we defined fROIs using the spatial working memory task (the *Hard* > *Easy* contrast), which has been shown to be effective in identifying the domain-general Multiple Demand network (Duncan, 2010; Fedorenko et al., 2013; Assem et al., 2020; Duncan et al., 2020) and examined their responses to the three tasks in our study.

Of the 27 language parcels in the extended set derived from the GSS analysis, 20 contained fROIs that reliably responded to the spatial working memory contrast, as estimated in an independent portion of the data (see **Figure S2.1**), with the exception of the right anterior temporal, medial frontal, medial anterior SFG, and left temporal pole, superior precuneus, precuneus / ACC, and BTLA. The fROIs that responded reliably to task demands showed no response to the language contrasts (or the responses were weak and not generalizable across modalities), which suggests that the language fROIs and these Multiple Demand fROIs are largely non-overlapping within these parcels. Only one MD fROI—within the left anterior temporal parcel—responded to both the reading-based and the auditory language contrasts (reading: $\beta = 0.20$, $p < .001$; auditory: $\beta = 0.47$, $p = .022$); however, in this fROI, the response to the working memory conditions is barely above baseline, and the *Hard* > *Easy* effect is small ($\beta = 0.07$), which suggests that the responses in this parcel are dominated by language processing, and even voxels that are selected to respond the most to the MD contrast inevitably show overlap with the language-responsive voxels.

Additionally, in the Harvard-Oxford Subcortical atlas (see **Figure S2.2.**), 10 of the 14 parcels contained fROIs that reliably responded to the spatial working memory contrast, including bilateral thalamus, caudate, putamen, and pallidum as well as the right hippocampus and accumbens. None of these fROIs responded reliably to the language conditions, which suggests that the language fROIs and these MD fROIs are non-overlapping within these parcels.

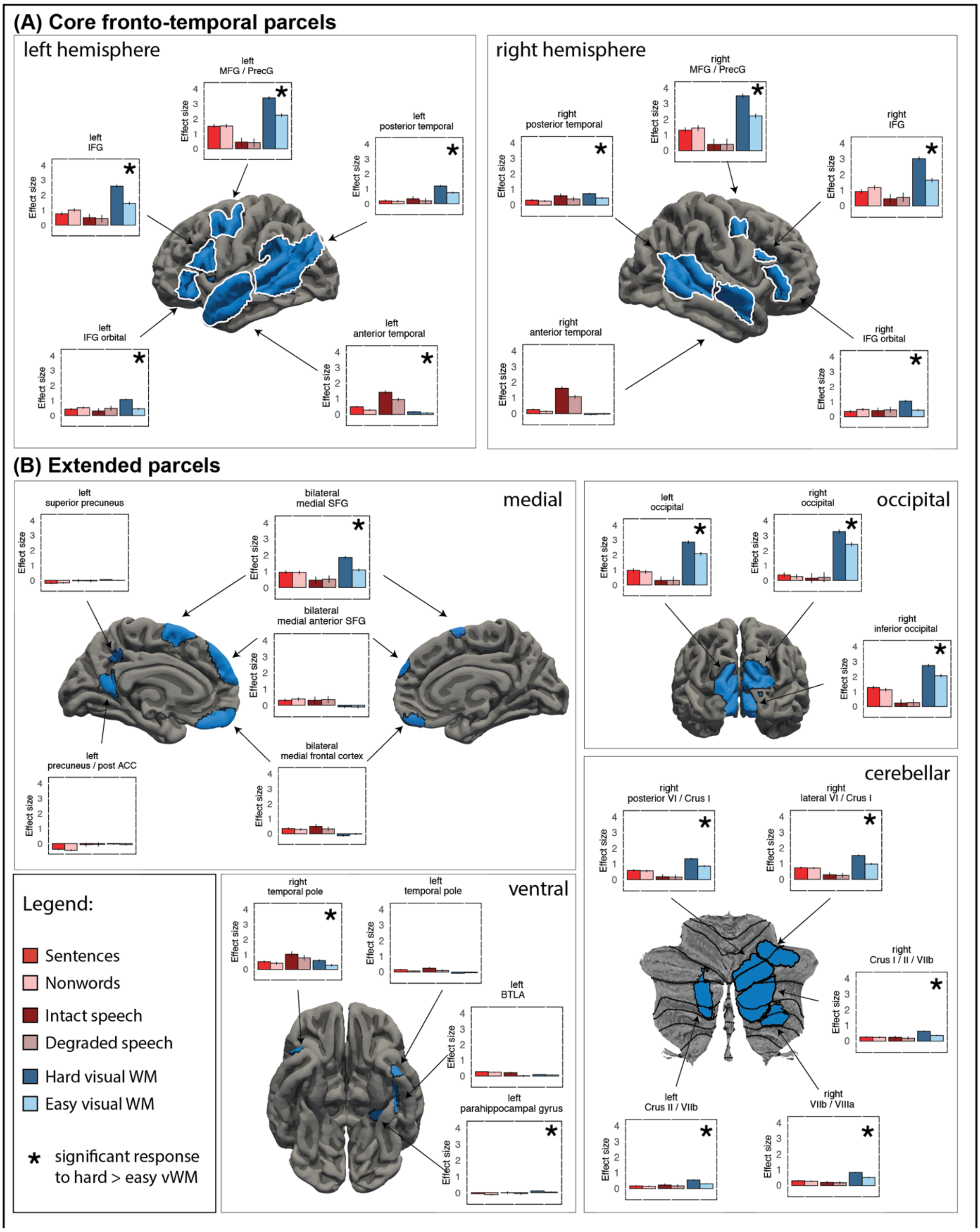


**Figure S2.1. Functional profiles of the MD fROIs within the new, extended set of language parcels.** Panel (A) shows the five core left-hemisphere regions and their right-hemisphere homotopes which are additionally highlighted with the white outlines. Panel (B) shows the extended set of parcels (as identified by the GSS procedure). Separate sub-panels show medial, occipital, ventral, and cerebellar parcels. The figure shows the surface projections of the cortical and cerebellar parcels (all the analyses were performed in the volume space, as described in Methods; the surface projections were created using FreeSurfer for the cortical and SUIT for the cerebellar parcels and are only used for visualization). For each region (defined based on individual activation maps, as described in Methods), we show responses to the reading-based and auditory language localizers (red bars; darker bars correspond to the critical conditions) and the non-linguistic spatial working memory task (blue bars; the darker bar corresponds to the harder condition). For the spatial working memory task, the responses were estimated using independent runs of the data. Parcels that contain fROIs showing a reliable response to the spatial WM contrast (i.e., *Hard* > *Easy*) are marked with a black asterisk in the upper right-hand corner.

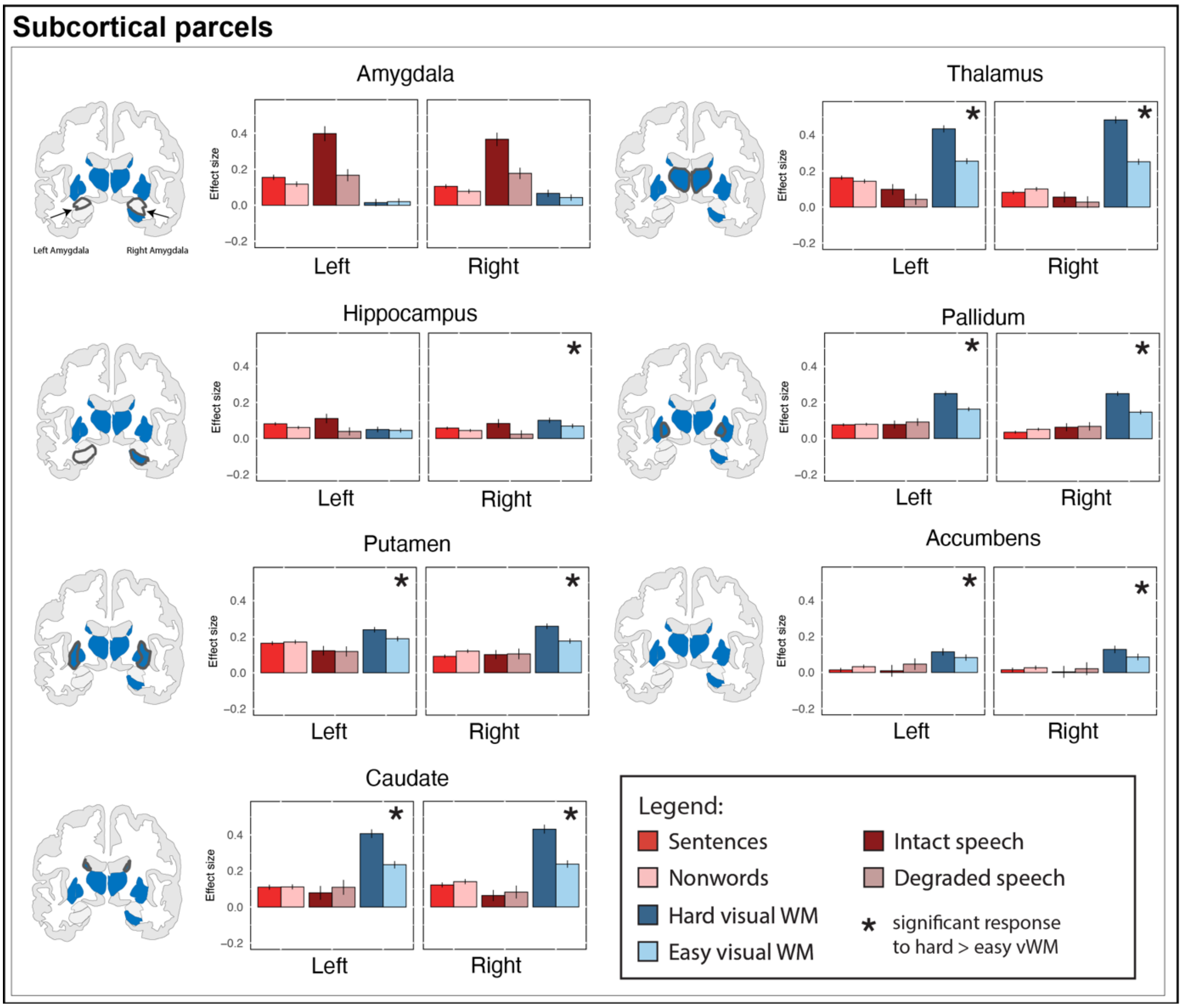


**Figure S2.2. Functional profiles of the MD fROIs within the parcels of the Harvard-Oxford Subcortical atlas.** For each region (defined based on individual activation maps; Methods), we show responses to the reading-based and auditory language localizers (red bars; darker bars correspond to the critical conditions) and the non-linguistic spatial working memory task (blue bars; the darker bar corresponds to the harder condition). For the non-linguistic spatial working memory task, the responses were estimated using independent runs of the data. Parcels that contain fROIs showing a reliable response to the spatial WM contrast (i.e., *Hard* > *Easy*) ) are marked with a black asterisk in the upper right-hand corner.

#### Section 3: The relationship between language activations and standard anatomical atlases.

To quantitatively compare the locations of the functional parcels (based on GSS parcellation) and the parcels from commonly used anatomical atlases, we calculated the percent of voxels in each atlas parcel that overlaps with any of the 27 functional parcels (e.g., if an entire atlas parcel falls within the boundaries of any of the functional parcels, the overlap will equal 100%, and if it falls fully outside the boundaries of the functional parcels, the overlap will be 0%). The results are presented in **Figure S3.1.A**. As evident from the figure, the general left-lateralized fronto-temporal topography is captured in all three parcellations, but coarse anatomical parcellations (such as DKT and HOC) create the appearance of a much more diffuse topography. One could argue however, that if you use individually defined functional ROIs, then it doesn’t much matter which parcels are used. However, some anatomical parcels are so large that they may encompass multiple functional areas. In other cases, a functional region may lie at an intersection of two macro-anatomical areas (e.g., straddling the middle frontal and precentral gyri). Both of these issues are illustrated in **Figure S3.1.B**. For these reasons, functional parcels, which are derived with knowledge of the functional topography of the relevant system, have advantages over anatomy-derived parcels.

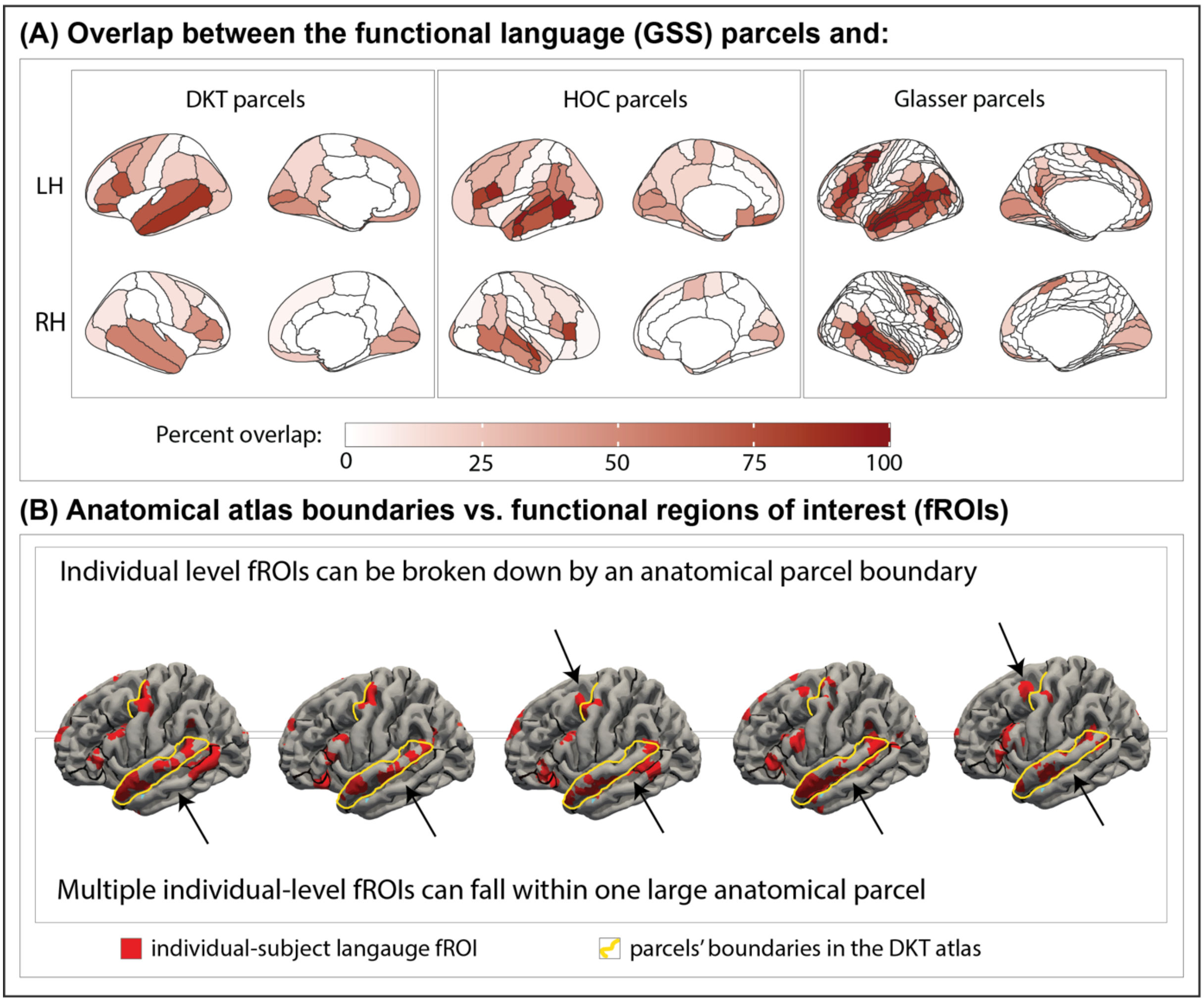

**Figure S3.1. The relationship between language activations and standard anatomical atlases.** Panel (A) presents the overlap between functional group-level language parcels and standard atlas parcels. For each parcel, the shade of red indicates the percentage of overlap with the functional parcels (darker shade = higher overlap). Panel (B) illustrates the two problems that can arise when using anatomical parcels from standard atlases (instead of functional parcels) to constrain the definition of individual fROIs. We are showing language activations defined as top 10% of most active voxels constrained by the group-level extended parcels set for five sample participants (red), with the yellow outlines highlighting two examples of parcels boundaries in the Desikan-Killiany-Tourville atlas. The top panel highlights examples of individual fROIs being broken up by a boundary between two atlas parcels (MFG and Precentral Gyrus); and the bottom panel highlights cases where a single parcel (STG) encompasses multiple fROIs (those that fall within the anterior temporal and the posterior temporal functional parcels).

For completeness, in **Figure S3.2**, we show the effect sizes for our three contrasts (the reading-based language localizer contrast, the auditory language localizer contrast, and the hard spatial working memory contrast) in the fROIs defined within all the parcels in the three atlases examined in this study (for details of the statistical tests, see **Table OSF 2** at https://osf.io/7594t/).

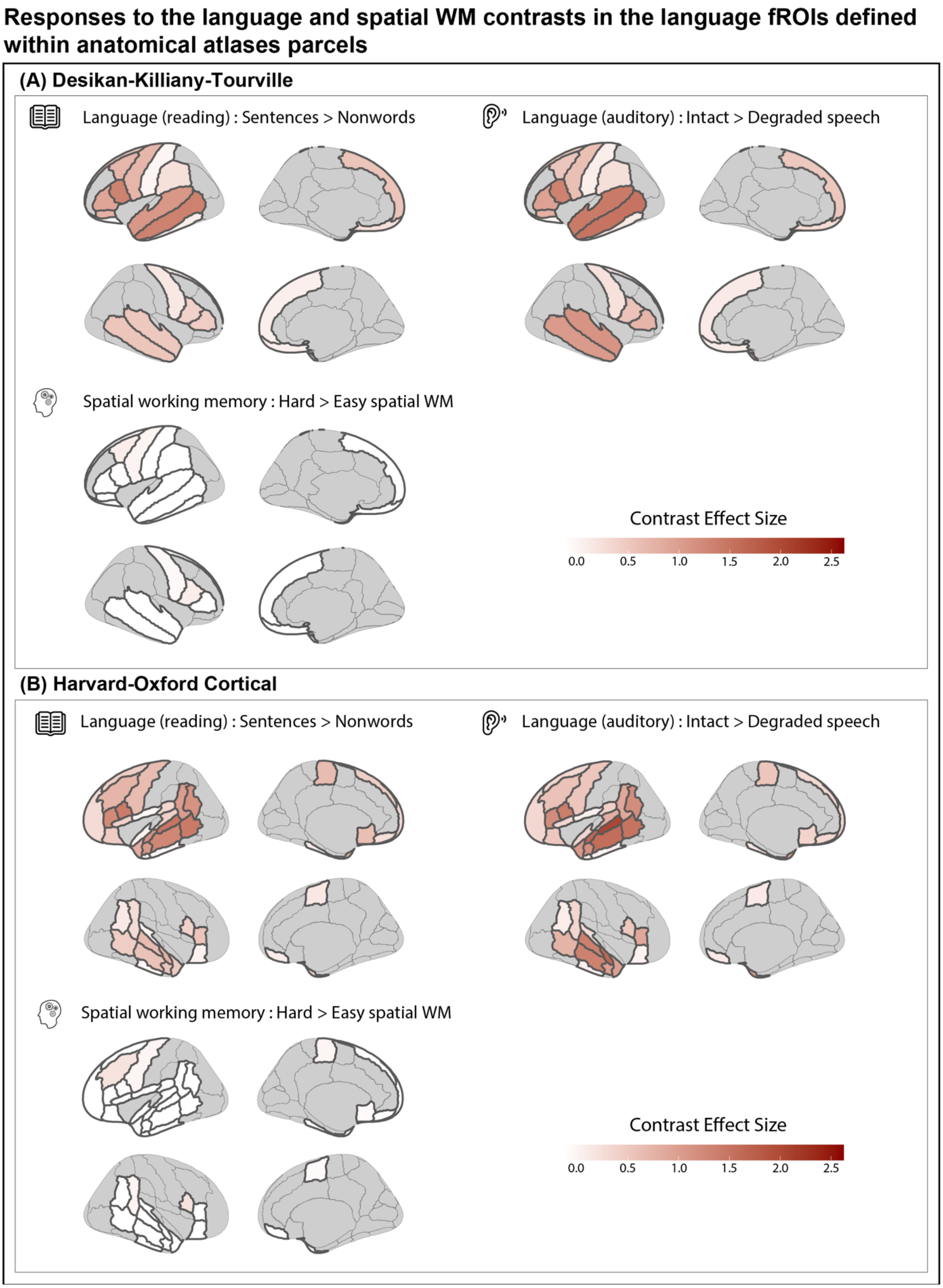
Responses to the language and spatial WM contrasts in the language fROIs defined within anatomical atlases parcels
(A) Desikan-Killiany-Tourville
Language (reading) : Sentences > Nonwords
Language (auditory) : Intact > Degraded speech
Spatial working memory : Hard > Easy spatial WM
Contrast Effect Size
0.0
0.5
1.0
1.5
2.0
2.5
(B) Harvard-Oxford Cortical
Language (reading) : Sentences > Nonwords
Language (auditory) : Intact > Degraded speech
Spatial working memory : Hard > Easy spatial WM
Contrast Effect Size
0.0
0.5
1.0
1.5
2.0
2.5

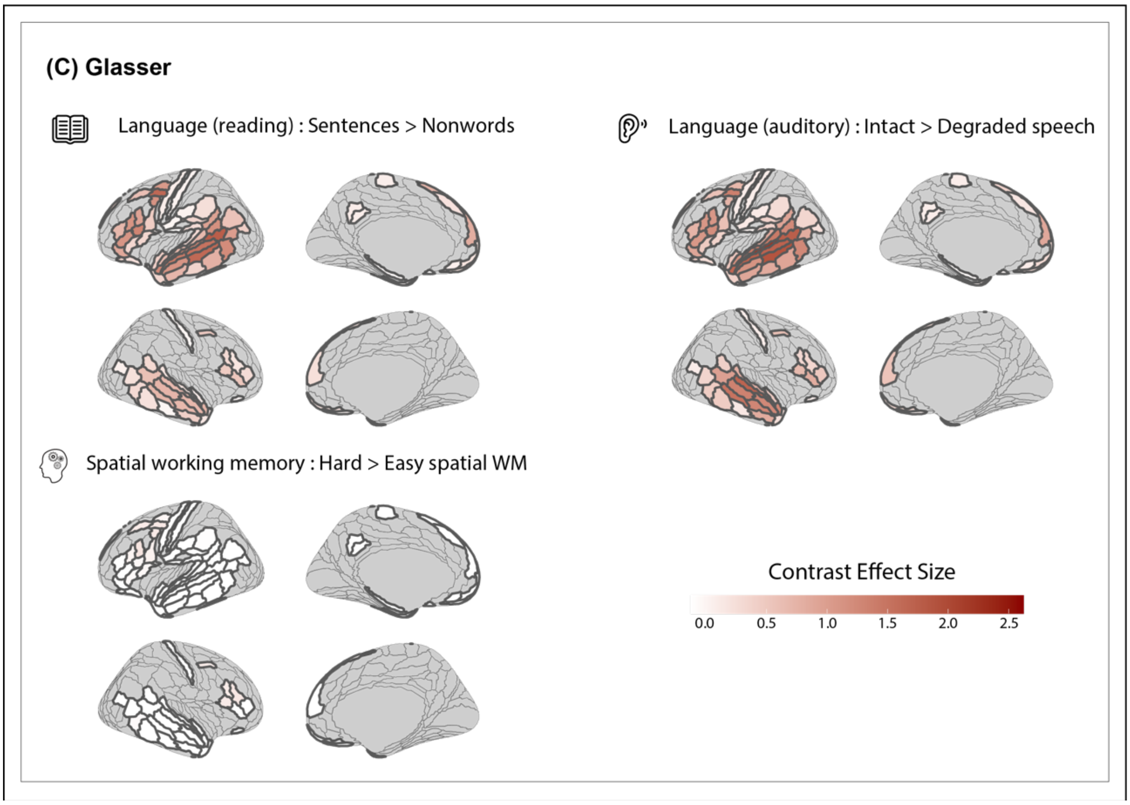


**Figure S3.2. Language responsiveness and selectivity of the language fROIs defined within the parcels in three standard anatomical atlases.** The three panels (A, B, and C) correspond to the three atlases (DKT: Klein & Tourville, 2012; HOC: Desikan et al., 2006; Glasser: Glasser et al., 2016, respectively). For each atlas and each contrast, we show 4 brain surfaces: left and right lateral and medial. For each contrast, the effect size is shown using the scale shown in each panel. Note that in these visualizations, we color the entire parcels, but the effects are estimated in the individually-defined fROIs (i.e., top 10% of the most language-responsive voxels in each parcel; Methods).

**Table S1. Statistical results for the key contrasts across the three paradigms (reading-based language localizer, auditory language localizer, and spatial working memory) in the language regions a) from the GSS analysis, and b) from the Harvard-Oxford Subcortical atlas.** The table presents estimates, standard errors, z-scores, and p-values for each condition (S, N, I, D, H, and E; see below for full condition names) relative to the fixation baseline, for each critical contrast (S>N, I>D, and H>E), and for the contrasts between the language conditions and the hard non-linguistic task condition (S>H and I>H), to evaluate language selectivity.
*S = Sentences; N = Nonwords, I = Intact Speech; D = Degraded Speech; H = Hard spatial WM; E = Easy spatial WM*

| Hemisphere | Label | Task task | Contrast | Effect Size | Std.Error | *z*-score | *p*-value |
|---|---|---|---|---|---|---|---|
| **Core cortical areas** | | | | | | | |
| left | IFG orbital | Reading Language Task | S | 1.52 | 0.02 | 65.55 | <.001 |
| | | | N | 0.44 | 0.02 | 18.98 | <.001 |
| | | | S>N | 1.08 | 0.02 | 61.78 | <.001 |
| | | | S>H | 2.07 | 0.03 | 63.79 | <.001 |
| | | Auditory Language Task | I | 1.74 | 0.08 | 22.60 | <.001 |
| | | | D | 0.71 | 0.08 | 9.24 | <.001 |
| | | | I>D | 1.03 | 0.1 | 10.25 | <.001 |

| | | | | | | | |
|---|---|---|---|---|---|---|---|
| | | | I>H | 2.3 | 0.08 | 28.78 | <.001 |
| | | Spatial WM Task | H | -0.55 | 0.04 | -15.44 | <.001 |
| | | | E | -0.34 | 0.04 | -9.51 | <.001 |
| | | | H>E | -0.21 | 0.04 | -5.11 | <.001 |
| left | IFG | Reading Language Task | S | 1.9 | 0.02 | 77.26 | <.001 |
| | | | N | 0.61 | 0.02 | 24.97 | <.001 |
| | | | S>N | 1.28 | 0.02 | 72.54 | <.001 |
| | | | S>H | 1.76 | 0.03 | 52.80 | <.001 |
| | | Auditory Language Task | I | 2.12 | 0.08 | 26.52 | <.001 |
| | | | D | 0.86 | 0.08 | 10.73 | <.001 |
| | | | I>D | 1.26 | 0.1 | 12.16 | <.001 |
| | | | I>H | 1.98 | 0.08 | 24.06 | <.001 |
| | | Spatial WM Task | H | 0.14 | 0.04 | 3.74 | <.001 |
| | | | E | 0.09 | 0.04 | 2.35 | .019 |
| | | | H>E | 0.05 | 0.04 | 1.22 | .224 |
| left | MFG / PrecG | Reading Language Task | S | 2.5 | 0.03 | 80.22 | <.001 |
| | | | N | 1.13 | 0.03 | 36.34 | <.001 |
| | | | S>N | 1.37 | 0.02 | 69.52 | <.001 |
| | | | S>H | 1.48 | 0.04 | 40.01 | <.001 |
| | | Auditory Language Task | I | 1.8 | 0.09 | 19.96 | <.001 |
| | | | D | 0.74 | 0.09 | 8.21 | <.001 |
| | | | I>D | 1.06 | 0.12 | 9.20 | <.001 |
| | | | I>H | 0.79 | 0.09 | 8.58 | <.001 |
| | | Spatial WM Task | H | 1.02 | 0.04 | 22.99 | <.001 |
| | | | E | 0.84 | 0.04 | 18.98 | <.001 |
| | | | H>E | 0.18 | 0.05 | 3.73 | <.001 |
| left | anterior temporal | Reading Language Task | S | 1.27 | 0.01 | 85.32 | <.001 |
| | | | N | 0.2 | 0.01 | 13.21 | <.001 |
| | | | S>N | 1.07 | 0.01 | 97.66 | <.001 |
| | | | S>H | 1.73 | 0.02 | 83.23 | <.001 |
| | | Auditory Language Task | I | 2.55 | 0.05 | 51.35 | <.001 |
| | | | D | 1.12 | 0.05 | 22.48 | <.001 |
| | | | I>D | 1.43 | 0.06 | 22.16 | <.001 |
| | | | I>H | 3.01 | 0.05 | 58.68 | <.001 |
| | | Spatial WM Task | H | -0.46 | 0.02 | -19.95 | <.001 |
| | | | E | -0.22 | 0.02 | -9.53 | <.001 |
| | | | H>E | -0.24 | 0.03 | -9.01 | <.001 |
| left | posterior temporal | Reading Language Task | S | 1.74 | 0.02 | 89.12 | <.001 |
| | | | N | 0.49 | 0.02 | 24.97 | <.001 |
| | | | S>N | 1.25 | 0.01 | 92.06 | <.001 |
| | | | S>H | 2.02 | 0.03 | 78.38 | <.001 |
| | | | I | 2.37 | 0.06 | 38.28 | <.001 |

| | | | | | | | |
|---|---|---|---|---|---|---|---|
| | | Auditory Language Task | D | 1 | 0.06 | 16.10 | <.001 |
| | | | I>D | 1.37 | 0.08 | 17.14 | <.001 |
| | | | I>H | 2.65 | 0.06 | 41.67 | <.001 |
| | | Spatial WM Task | H | -0.28 | 0.03 | -9.54 | <.001 |
| | | | E | -0.09 | 0.03 | -3.00 | .003 |
| | | | H>E | -0.19 | 0.03 | -5.80 | <.001 |
| right | IFG orbital | Reading Language Task | S | 0.96 | 0.02 | 43.24 | <.001 |
| | | | N | 0.26 | 0.02 | 11.66 | <.001 |
| | | | S>N | 0.7 | 0.02 | 37.65 | <.001 |
| | | | S>H | 1.31 | 0.03 | 39.74 | <.001 |
| | | Auditory Language Task | I | 1.61 | 0.08 | 20.76 | <.001 |
| | | | D | 0.76 | 0.08 | 9.81 | <.001 |
| | | | I>D | 0.85 | 0.1 | 8.34 | <.001 |
| | | | I>H | 1.96 | 0.08 | 24.39 | <.001 |
| | | Spatial WM Task | H | -0.36 | 0.04 | -10.18 | <.001 |
| | | | E | -0.24 | 0.04 | -6.72 | <.001 |
| | | | H>E | -0.12 | 0.04 | -2.90 | .004 |
| right | IFG | Reading Language Task | S | 1.07 | 0.03 | 41.88 | <.001 |
| | | | N | 0.39 | 0.03 | 15.34 | <.001 |
| | | | S>N | 0.68 | 0.02 | 32.90 | <.001 |
| | | | S>H | 0.71 | 0.04 | 19.68 | <.001 |
| | | Auditory Language Task | I | 1.58 | 0.08 | 18.78 | <.001 |
| | | | D | 0.86 | 0.08 | 10.17 | <.001 |
| | | | I>D | 0.73 | 0.11 | 6.61 | <.001 |
| | | | I>H | 1.22 | 0.09 | 14.03 | <.001 |
| | | Spatial WM Task | H | 0.36 | 0.04 | 9.25 | <.001 |
| | | | E | 0.17 | 0.04 | 4.48 | <.001 |
| | | | H>E | 0.18 | 0.05 | 4.09 | <.001 |
| right | MFG /PrecG | Reading Language Task | S | 1.36 | 0.03 | 40.82 | <.001 |
| | | | N | 0.72 | 0.03 | 21.63 | <.001 |
| | | | S>N | 0.64 | 0.02 | 26.72 | <.001 |
| | | | S>H | 0.67 | 0.04 | 15.77 | <.001 |
| | | Auditory Language Task | I | 1.78 | 0.1 | 17.77 | <.001 |
| | | | D | 1.14 | 0.1 | 11.42 | <.001 |
| | | | I>D | 0.64 | 0.13 | 4.93 | <.001 |
| | | | I>H | 1.08 | 0.1 | 10.55 | <.001 |
| | | Spatial WM Task | H | 0.7 | 0.05 | 14.58 | <.001 |
| | | | E | 0.57 | 0.05 | 11.93 | <.001 |
| | | | H>E | 0.13 | 0.05 | 2.39 | .017 |
| right | anterior temporal | Reading Language Task | S | 0.88 | 0.02 | 50.87 | <.001 |
| | | | N | 0.07 | 0.02 | 4.22 | <.001 |
| | | | S>N | 0.81 | 0.01 | 64.31 | <.001 |

| | | | | | | | |
|---|---|---|---|---|---|---|---|
| | | | S>H | 1.44 | 0.02 | 61.41 | <.001 |
| | | Auditory Language Task | I | 2.8 | 0.06 | 50.11 | <.001 |
| | | | D | 1.33 | 0.06 | 23.87 | <.001 |
| | | | I>D | 1.47 | 0.07 | 20.23 | <.001 |
| | | | I>H | 3.35 | 0.06 | 58.30 | <.001 |
| | | Spatial WM Task | H | -0.55 | 0.03 | -21.11 | <.001 |
| | | | E | -0.29 | 0.03 | -11.00 | <.001 |
| | | | H>E | -0.26 | 0.03 | -8.88 | <.001 |
| right | posterior temporal | Reading Language Task | S | 1.12 | 0.02 | 57.68 | <.001 |
| | | | N | 0.39 | 0.02 | 19.93 | <.001 |
| | | | S>N | 0.73 | 0.01 | 54.73 | <.001 |
| | | | S>H | 1.28 | 0.03 | 51.00 | <.001 |
| | | Auditory Language Task | I | 2.44 | 0.06 | 40.59 | <.001 |
| | | | D | 1.33 | 0.06 | 22.15 | <.001 |
| | | | I>D | 1.11 | 0.08 | 14.30 | <.001 |
| | | | I>H | 2.6 | 0.06 | 42.14 | <.001 |
| | | Spatial WM Task | H | -0.15 | 0.03 | -5.41 | <.001 |
| | | | E | -0.02 | 0.03 | -0.86 | .389 |
| | | | H>E | -0.13 | 0.03 | -4.08 | <.001 |
| **Extended cortical areas** | | | | | | | |
| left | temporal pole | Reading Language Task | S | 0.37 | 0.01 | 32.26 | <.001 |
| | | | N | 0 | 0.01 | 0.20 | .838 |
| | | | S>N | 0.37 | 0.01 | 36.88 | <.001 |
| | | | S>H | 0.61 | 0.02 | 36.85 | <.001 |
| | | Auditory Language Task | I | 0.41 | 0.04 | 10.71 | <.001 |
| | | | D | 0.12 | 0.04 | 3.11 | .002 |
| | | | I>D | 0.29 | 0.05 | 5.81 | <.001 |
| | | | I>H | 0.65 | 0.04 | 16.33 | <.001 |
| | | Spatial WM Task | H | -0.24 | 0.02 | -13.69 | <.001 |
| | | | E | -0.12 | 0.02 | -6.67 | <.001 |
| | | | H>E | -0.12 | 0.02 | -5.95 | <.001 |
| left | BTLA | Reading Language Task | S | 0.38 | 0.01 | 38.82 | <.001 |
| | | | N | 0.1 | 0.01 | 10.28 | <.001 |
| | | | S>N | 0.28 | 0.01 | 32.86 | <.001 |
| | | | S>H | 0.45 | 0.01 | 33.13 | <.001 |
| | | Auditory Language Task | I | 0.32 | 0.03 | 10.11 | <.001 |
| | | | D | -0.03 | 0.03 | -1.04 | .299 |
| | | | I>D | 0.35 | 0.04 | 8.55 | <.001 |
| | | | I>H | 0.39 | 0.03 | 12.04 | <.001 |
| | | Spatial WM Task | H | -0.07 | 0.01 | -5.10 | <.001 |
| | | | E | -0.01 | 0.01 | -1.02 | .309 |

| | | | | | | | |
|---|---|---|---|---|---|---|---|
| | | | H>E | -0.06 | 0.02 | -3.50 | <.001 |
| left | parahippocampal gyrus | Reading Language Task | S | 0.03 | 0.01 | 3.03 | .002 |
| | | | N | -0.19 | 0.01 | -18.73 | <.001 |
| | | | S>N | 0.22 | 0.01 | 23.88 | <.001 |
| | | | S>H | 0.2 | 0.01 | 13.54 | <.001 |
| | | Auditory Language Task | I | 0.02 | 0.03 | 0.62 | .534 |
| | | | D | -0.08 | 0.03 | -2.41 | .016 |
| | | | I>D | 0.1 | 0.04 | 2.31 | .021 |
| | | | I>H | 0.19 | 0.03 | 5.43 | <.001 |
| | | Spatial WM Task | H | -0.17 | 0.01 | -11.03 | <.001 |
| | | | E | -0.11 | 0.01 | -7.16 | <.001 |
| | | | H>E | -0.06 | 0.02 | -3.27 | .001 |
| left | superior precuneus | Reading Language Task | S | 0.22 | 0.01 | 17.35 | <.001 |
| | | | N | -0.16 | 0.01 | -13.21 | <.001 |
| | | | S>N | 0.38 | 0.01 | 32.63 | <.001 |
| | | | S>H | 0.81 | 0.02 | 41.19 | <.001 |
| | | Auditory Language Task | I | 0.51 | 0.05 | 11.29 | <.001 |
| | | | D | 0.21 | 0.05 | 4.54 | <.001 |
| | | | I>D | 0.31 | 0.06 | 5.10 | <.001 |
| | | | I>H | 1.11 | 0.05 | 23.36 | <.001 |
| | | Spatial WM Task | H | -0.6 | 0.02 | -29.64 | <.001 |
| | | | E | -0.34 | 0.02 | -17.03 | <.001 |
| | | | H>E | -0.25 | 0.02 | -10.28 | <.001 |
| left | precuneus / post CC | Reading Language Task | S | -0.11 | 0.02 | -6.80 | <.001 |
| | | | N | -0.35 | 0.02 | -22.24 | <.001 |
| | | | S>N | 0.24 | 0.01 | 17.96 | <.001 |
| | | | S>H | 0.46 | 0.02 | 20.75 | <.001 |
| | | Auditory Language Task | I | 0.07 | 0.05 | 1.39 | .163 |
| | | | D | -0.07 | 0.05 | -1.37 | .172 |
| | | | I>D | 0.14 | 0.07 | 2.11 | .035 |
| | | | I>H | 0.64 | 0.05 | 12.04 | <.001 |
| | | Spatial WM Task | H | -0.57 | 0.02 | -24.13 | <.001 |
| | | | E | -0.34 | 0.02 | -14.27 | <.001 |
| | | | H>E | -0.23 | 0.03 | -8.43 | <.001 |
| left | occipital | Reading Language Task | S | 0.12 | 0.04 | 3.08 | .002 |
| | | | N | -0.26 | 0.04 | -6.98 | <.001 |
| | | | S>N | 0.38 | 0.02 | 17.10 | <.001 |
| | | | S>H | -1.16 | 0.04 | -29.89 | <.001 |
| | | Auditory Language Task | I | -0.23 | 0.1 | -2.46 | .014 |
| | | | D | -0.25 | 0.1 | -2.66 | .008 |
| | | | I>D | 0.02 | 0.12 | 0.16 | .874 |
| | | | I>H | -1.51 | 0.09 | -15.94 | <.001 |

| | | | | | | | |
|---|---|---|---|---|---|---|---|
| | | Spatial WM Task | H | 1.28 | 0.05 | 26.02 | <.001 |
| | | | E | 0.94 | 0.05 | 19.01 | <.001 |
| | | | H>E | 0.34 | 0.05 | 7.02 | <.001 |
| right | temporal pole | Reading Language Task | S | 0.91 | 0.02 | 46.08 | <.001 |
| | | | N | 0.15 | 0.02 | 7.48 | <.001 |
| | | | S>N | 0.76 | 0.02 | 47.71 | <.001 |
| | | | S>H | 1.14 | 0.03 | 39.35 | <.001 |
| | | Auditory Language Task | I | 2.17 | 0.07 | 31.78 | <.001 |
| | | | D | 0.92 | 0.07 | 13.51 | <.001 |
| | | | I>D | 1.25 | 0.09 | 13.95 | <.001 |
| | | | I>H | 2.4 | 0.07 | 33.88 | <.001 |
| | | Spatial WM Task | H | -0.23 | 0.03 | -7.44 | <.001 |
| | | | E | -0.04 | 0.03 | -1.39 | .165 |
| | | | H>E | -0.19 | 0.04 | -5.12 | <.001 |
| right | occipital | Reading Language Task | S | -0.19 | 0.04 | -4.85 | <.001 |
| | | | N | -0.57 | 0.04 | -14.41 | <.001 |
| | | | S>N | 0.38 | 0.03 | 14.83 | <.001 |
| | | | S>H | -1.48 | 0.04 | -33.88 | <.001 |
| | | Auditory Language Task | I | -0.09 | 0.1 | -0.87 | .387 |
| | | | D | -0.07 | 0.1 | -0.71 | .478 |
| | | | I>D | -0.02 | 0.13 | -0.12 | .902 |
| | | | I>H | -1.37 | 0.11 | -13.06 | <.001 |
| | | Spatial WM Task | H | 1.28 | 0.05 | 24.26 | <.001 |
| | | | E | 0.94 | 0.05 | 17.70 | <.001 |
| | | | H>E | 0.35 | 0.05 | 6.38 | <.001 |
| right | inferior occipital | Reading Language Task | S | 0.68 | 0.03 | 20.31 | <.001 |
| | | | N | 0.31 | 0.03 | 9.15 | <.001 |
| | | | S>N | 0.38 | 0.02 | 16.59 | <.001 |
| | | | S>H | -1.26 | 0.04 | -31.70 | <.001 |
| | | Auditory Language Task | I | 0.05 | 0.1 | 0.56 | .576 |
| | | | D | 0.02 | 0.1 | 0.17 | .864 |
| | | | I>D | 0.04 | 0.12 | 0.30 | .762 |
| | | | I>H | -1.89 | 0.1 | -19.57 | <.001 |
| | | Spatial WM Task | H | 1.94 | 0.05 | 41.58 | <.001 |
| | | | E | 1.55 | 0.05 | 33.17 | <.001 |
| | | | H>E | 0.39 | 0.05 | 7.87 | <.001 |
| bilateral | ventromedial frontal cortex | Reading Language Task | S | 0.61 | 0.02 | 28.64 | <.001 |
| | | | N | 0.16 | 0.02 | 7.70 | <.001 |
| | | | S>N | 0.45 | 0.02 | 23.69 | <.001 |
| | | | S>H | 1.44 | 0.03 | 44.62 | <.001 |
| | | Auditory Language Task | I | 0.8 | 0.07 | 10.76 | <.001 |
| | | | D | 0.36 | 0.07 | 4.83 | <.001 |

| | | | | | | | |
|---|---|---|---|---|---|---|---|
| | | | I>D | 0.44 | 0.1 | 4.51 | <.001 |
| | | | I>H | 1.63 | 0.08 | 20.98 | <.001 |
| | | Spatial WM Task | H | -0.83 | 0.03 | -24.66 | <.001 |
| | | | E | -0.35 | 0.03 | -10.27 | <.001 |
| | | | H>E | -0.48 | 0.04 | -11.96 | <.001 |
| bilateral | medial anterior SFG | Reading Language Task | S | 1.19 | 0.03 | 40.47 | <.001 |
| | | | N | 0.12 | 0.03 | 4.04 | <.001 |
| | | | S>N | 1.07 | 0.02 | 43.79 | <.001 |
| | | | S>H | 2.5 | 0.04 | 56.02 | <.001 |
| | | Auditory Language Task | I | 1.4 | 0.11 | 13.31 | <.001 |
| | | | D | 0.43 | 0.11 | 4.07 | <.001 |
| | | | I>D | 0.97 | 0.14 | 7.02 | <.001 |
| | | | I>H | 2.71 | 0.11 | 24.78 | <.001 |
| | | Spatial WM Task | H | -1.31 | 0.05 | -27.72 | <.001 |
| | | | E | -0.64 | 0.05 | -13.48 | <.001 |
| | | | H>E | -0.68 | 0.06 | -11.86 | <.001 |
| bilateral | medial SFG | Reading Language Task | S | 1.7 | 0.03 | 60.75 | <.001 |
| | | | N | 0.74 | 0.03 | 26.59 | <.001 |
| | | | S>N | 0.96 | 0.02 | 51.10 | <.001 |
| | | | S>H | 1.21 | 0.03 | 34.76 | <.001 |
| | | Auditory Language Task | I | 1.09 | 0.08 | 13.00 | <.001 |
| | | | D | 0.52 | 0.08 | 6.18 | <.001 |
| | | | I>D | 0.57 | 0.11 | 5.30 | <.001 |
| | | | I>H | 0.61 | 0.09 | 7.06 | <.001 |
| | | Spatial WM Task | H | 0.49 | 0.04 | 12.03 | <.001 |
| | | | E | 0.41 | 0.04 | 10.22 | <.001 |
| | | | H>E | 0.07 | 0.04 | 1.65 | .099 |
| **Cerebellar areas** | | | | | | | |
| left | Crus I / II / VIIb | Reading Language Task | S | 0.28 | 0.01 | 28.26 | <.001 |
| | | | N | 0.07 | 0.01 | 6.96 | <.001 |
| | | | S>N | 0.21 | 0.01 | 23.50 | <.001 |
| | | | S>H | 0.24 | 0.01 | 16.36 | <.001 |
| | | Auditory Language Task | I | 0.44 | 0.03 | 12.82 | <.001 |
| | | | D | 0.24 | 0.03 | 7.07 | <.001 |
| | | | I>D | 0.2 | 0.04 | 4.37 | <.001 |
| | | | I>H | 0.4 | 0.04 | 11.22 | <.001 |
| | | Spatial WM Task | H | 0.04 | 0.02 | 2.52 | .012 |
| | | | E | 0.03 | 0.02 | 2.19 | .028 |
| | | | H>E | 0.01 | 0.02 | 0.27 | .785 |
| right | Crus I / II / VIIb | Reading Language Task | S | 0.56 | 0.01 | 53.89 | <.001 |
| | | | N | 0.14 | 0.01 | 13.32 | <.001 |

| | | | | | | | |
|---|---|---|---|---|---|---|---|
| | | | S>N | 0.42 | 0.01 | 49.26 | <.001 |
| | | | S>H | 0.63 | 0.02 | 40.98 | <.001 |
| | | Auditory Language Task | I | 0.6 | 0.04 | 16.66 | <.001 |
| | | | D | 0.31 | 0.04 | 8.66 | <.001 |
| | | | I>D | 0.29 | 0.05 | 6.10 | <.001 |
| | | | I>H | 0.68 | 0.04 | 17.99 | <.001 |
| | | Spatial WM Task | H | -0.07 | 0.02 | -4.52 | <.001 |
| | | | E | -0.01 | 0.02 | -0.73 | .466 |
| | | | H>E | -0.06 | 0.02 | -3.18 | .001 |
| right | VIIb / VIIIa | Reading Language Task | S | 0.73 | 0.01 | 58.47 | <.001 |
| | | | N | 0.34 | 0.01 | 27.48 | <.001 |
| | | | S>N | 0.39 | 0.01 | 46.10 | <.001 |
| | | | S>H | 0.45 | 0.01 | 30.72 | <.001 |
| | | Auditory Language Task | I | 0.66 | 0.04 | 18.65 | <.001 |
| | | | D | 0.41 | 0.04 | 11.74 | <.001 |
| | | | I>D | 0.24 | 0.04 | 5.42 | <.001 |
| | | | I>H | 0.38 | 0.04 | 10.61 | <.001 |
| | | Spatial WM Task | H | 0.28 | 0.02 | 16.05 | <.001 |
| | | | E | 0.2 | 0.02 | 11.81 | <.001 |
| | | | H>E | 0.07 | 0.02 | 3.97 | <.001 |
| right | posterior VI / Crus I | Reading Language Task | S | 0.57 | 0.02 | 37.50 | <.001 |
| | | | N | 0.23 | 0.02 | 15.22 | <.001 |
| | | | S>N | 0.34 | 0.01 | 28.87 | <.001 |
| | | | S>H | 0.03 | 0.02 | 1.32 | .187 |
| | | Auditory Language Task | I | 0.39 | 0.05 | 8.10 | <.001 |
| | | | D | 0.22 | 0.05 | 4.51 | <.001 |
| | | | I>D | 0.17 | 0.06 | 2.76 | .006 |
| | | | I>H | -0.16 | 0.05 | -3.17 | .002 |
| | | Spatial WM Task | H | 0.55 | 0.02 | 24.22 | <.001 |
| | | | E | 0.41 | 0.02 | 18.07 | <.001 |
| | | | H>E | 0.14 | 0.03 | 5.42 | <.001 |
| right | lateral VI | Reading Language Task | S | 0.82 | 0.02 | 48.38 | <.001 |
| | | | N | 0.5 | 0.02 | 29.29 | <.001 |
| | | | S>N | 0.32 | 0.01 | 28.28 | <.001 |
| | | | S>H | -0.02 | 0.02 | -1.21 | .227 |
| | | Auditory Language Task | I | 0.52 | 0.05 | 11.15 | <.001 |
| | | | D | 0.33 | 0.05 | 7.14 | <.001 |
| | | | I>D | 0.19 | 0.06 | 3.16 | .002 |
| | | | I>H | -0.33 | 0.05 | -6.95 | <.001 |
| | | Spatial WM Task | H | 0.84 | 0.02 | 36.60 | <.001 |
| | | | E | 0.6 | 0.02 | 26.09 | <.001 |
| | | | H>E | 0.24 | 0.02 | 9.99 | <.001 |

| | | | | | | | |
|---|---|---|---|---|---|---|---|
| **Subcortical areas** | | | | | | | |
| left | thalamus | Reading Language Task | S | 0.16 | 0.01 | 19.75 | <.001 |
| | | | N | 0.12 | 0.01 | 14.76 | <.001 |
| | | | S>N | 0.04 | 0.01 | 4.69 | <.001 |
| | | | S>H | -0.1 | 0.01 | -7.91 | <.001 |
| | | Auditory Language Task | I | 0.15 | 0.03 | 5.49 | <.001 |
| | | | D | 0.06 | 0.03 | 2.02 | .043 |
| | | | I>D | 0.09 | 0.04 | 2.71 | .007 |
| | | | I>H | -0.11 | 0.03 | -3.79 | <.001 |
| | | Spatial WM Task | H | 0.26 | 0.01 | 21.95 | <.001 |
| | | | E | 0.16 | 0.01 | 13.88 | <.001 |
| | | | H>E | 0.09 | 0.01 | 6.57 | <.001 |
| left | caudate | Reading Language Task | S | 0.14 | 0.01 | 13.04 | <.001 |
| | | | N | 0.09 | 0.01 | 8.55 | <.001 |
| | | | S>N | 0.05 | 0.01 | 4.32 | <.001 |
| | | | S>H | -0.16 | 0.02 | -10.51 | <.001 |
| | | Auditory Language Task | I | 0.05 | 0.03 | 1.45 | .148 |
| | | | D | 0.06 | 0.03 | 1.70 | .089 |
| | | | I>D | -0.01 | 0.04 | -0.20 | .842 |
| | | | I>H | -0.25 | 0.04 | -6.96 | <.001 |
| | | Spatial WM Task | H | 0.3 | 0.01 | 20.01 | <.001 |
| | | | E | 0.2 | 0.01 | 13.63 | <.001 |
| | | | H>E | 0.1 | 0.02 | 5.27 | <.001 |
| left | putamen | Reading Language Task | S | 0.18 | 0.01 | 22.78 | <.001 |
| | | | N | 0.16 | 0.01 | 19.43 | <.001 |
| | | | S>N | 0.03 | 0.01 | 3.41 | .001 |
| | | | S>H | 0 | 0.01 | -0.42 | .671 |
| | | Auditory Language Task | I | 0.14 | 0.03 | 5.42 | <.001 |
| | | | D | 0.09 | 0.03 | 3.56 | <.001 |
| | | | I>D | 0.05 | 0.03 | 1.48 | .139 |
| | | | I>H | -0.05 | 0.03 | -1.90 | .058 |
| | | Spatial WM Task | H | 0.19 | 0.01 | 16.79 | <.001 |
| | | | E | 0.18 | 0.01 | 16.47 | <.001 |
| | | | H>E | 0 | 0.01 | 0.27 | .786 |
| left | pallidum | Reading Language Task | S | 0.12 | 0.01 | 17.05 | <.001 |
| | | | N | 0.09 | 0.01 | 13.92 | <.001 |
| | | | S>N | 0.02 | 0.01 | 3.13 | .002 |
| | | | S>H | -0.07 | 0.01 | -7.28 | <.001 |
| | | Auditory Language Task | I | 0.1 | 0.02 | 4.46 | <.001 |
| | | | D | 0.07 | 0.02 | 3.39 | .001 |
| | | | I>D | 0.02 | 0.03 | 0.85 | .395 |

| | | | | | | | |
|---|---|---|---|---|---|---|---|
| | | | I>H | -0.09 | 0.02 | -3.97 | <.001 |
| | | Spatial WM Task | H | 0.19 | 0.01 | 19.56 | <.001 |
| | | | E | 0.15 | 0.01 | 15.28 | <.001 |
| | | | H>E | 0.04 | 0.01 | 3.60 | <.001 |
| left | hippocampus | Reading Language Task | S | 0.14 | 0.01 | 17.76 | <.001 |
| | | | N | 0.05 | 0.01 | 6.13 | <.001 |
| | | | S>N | 0.09 | 0.01 | 10.80 | <.001 |
| | | | S>H | 0.32 | 0.01 | 27.23 | <.001 |
| | | Auditory Language Task | I | 0.24 | 0.03 | 9.06 | <.001 |
| | | | D | 0.08 | 0.03 | 3.15 | .002 |
| | | | I>D | 0.15 | 0.03 | 4.60 | <.001 |
| | | | I>H | 0.42 | 0.03 | 15.49 | <.001 |
| | | Spatial WM Task | H | -0.18 | 0.01 | -16.48 | <.001 |
| | | | E | -0.08 | 0.01 | -7.01 | <.001 |
| | | | H>E | -0.11 | 0.01 | -7.66 | <.001 |
| left | amygdala | Reading Language Task | S | 0.19 | 0.01 | 18.61 | <.001 |
| | | | N | 0.11 | 0.01 | 10.73 | <.001 |
| | | | S>N | 0.08 | 0.01 | 7.50 | <.001 |
| | | | S>H | 0.33 | 0.01 | 21.75 | <.001 |
| | | Auditory Language Task | I | 0.54 | 0.03 | 16.06 | <.001 |
| | | | D | 0.23 | 0.03 | 6.90 | <.001 |
| | | | I>D | 0.31 | 0.04 | 7.17 | <.001 |
| | | | I>H | 0.67 | 0.03 | 19.51 | <.001 |
| | | Spatial WM Task | H | -0.14 | 0.01 | -9.69 | <.001 |
| | | | E | -0.05 | 0.01 | -3.75 | <.001 |
| | | | H>E | -0.09 | 0.02 | -4.88 | <.001 |
| left | accumbens | Reading Language Task | S | 0.02 | 0.01 | 2.30 | .022 |
| | | | N | 0.02 | 0.01 | 1.85 | .064 |
| | | | S>N | 0 | 0.01 | 0.41 | .682 |
| | | | S>H | -0.03 | 0.01 | -1.83 | .068 |
| | | Auditory Language Task | I | -0.02 | 0.03 | -0.54 | .587 |
| | | | D | -0.03 | 0.03 | -1.09 | .276 |
| | | | I>D | 0.02 | 0.04 | 0.43 | .671 |
| | | | I>H | -0.06 | 0.03 | -1.96 | .050 |
| | | Spatial WM Task | H | 0.05 | 0.01 | 3.50 | <.001 |
| | | | E | 0.06 | 0.01 | 4.23 | <.001 |
| | | | H>E | -0.01 | 0.02 | -0.58 | .559 |
| right | thalamus | Reading Language Task | S | 0.07 | 0.01 | 9.67 | <.001 |
| | | | N | 0.07 | 0.01 | 8.91 | <.001 |
| | | | S>N | 0.01 | 0.01 | 0.71 | .477 |
| | | | S>H | -0.2 | 0.01 | -17.57 | <.001 |
| | | | I | 0.04 | 0.02 | 1.59 | .111 |

| | | | | | | | |
|---|---|---|---|---|---|---|---|
| | | Auditory Language Task | D | 0.01 | 0.02 | 0.43 | .665 |
| | | | I>D | 0.03 | 0.03 | 0.91 | .365 |
| | | | I>H | -0.23 | 0.03 | -8.84 | <.001 |
| | | Spatial WM Task | H | 0.27 | 0.01 | 25.12 | <.001 |
| | | | E | 0.14 | 0.01 | 13.35 | <.001 |
| | | | H>E | 0.13 | 0.01 | 9.58 | <.001 |
| right | caudate | Reading Language Task | S | 0.11 | 0.01 | 11.58 | <.001 |
| | | | N | 0.1 | 0.01 | 10.26 | <.001 |
| | | | S>N | 0.01 | 0.01 | 1.30 | .195 |
| | | | S>H | -0.15 | 0.01 | -10.83 | <.001 |
| | | Auditory Language Task | I | 0 | 0.03 | 0.12 | .905 |
| | | | D | -0.02 | 0.03 | -0.54 | .592 |
| | | | I>D | 0.02 | 0.04 | 0.52 | .605 |
| | | | I>H | -0.26 | 0.03 | -8.00 | <.001 |
| | | Spatial WM Task | H | 0.26 | 0.01 | 19.31 | <.001 |
| | | | E | 0.17 | 0.01 | 12.74 | <.001 |
| | | | H>E | 0.09 | 0.02 | 5.46 | <.001 |
| right | putamen | Reading Language Task | S | 0.08 | 0.01 | 9.96 | <.001 |
| | | | N | 0.08 | 0.01 | 10.78 | <.001 |
| | | | S>N | -0.01 | 0.01 | -0.79 | .429 |
| | | | S>H | -0.09 | 0.01 | -8.08 | <.001 |
| | | Auditory Language Task | I | 0.08 | 0.03 | 3.29 | .001 |
| | | | D | 0.06 | 0.03 | 2.51 | .012 |
| | | | I>D | 0.02 | 0.03 | 0.62 | .535 |
| | | | I>H | -0.08 | 0.03 | -3.24 | .001 |
| | | Spatial WM Task | H | 0.17 | 0.01 | 15.34 | <.001 |
| | | | E | 0.15 | 0.01 | 13.72 | <.001 |
| | | | H>E | 0.02 | 0.01 | 1.34 | .179 |
| right | pallidum | Reading Language Task | S | 0.04 | 0.01 | 6.13 | <.001 |
| | | | N | 0.04 | 0.01 | 5.85 | <.001 |
| | | | S>N | 0 | 0.01 | 0.28 | .778 |
| | | | S>H | -0.13 | 0.01 | -15.08 | <.001 |
| | | Auditory Language Task | I | 0.06 | 0.02 | 2.79 | .005 |
| | | | D | 0.04 | 0.02 | 2.18 | .029 |
| | | | I>D | 0.01 | 0.03 | 0.48 | .631 |
| | | | I>H | -0.12 | 0.02 | -5.66 | <.001 |
| | | Spatial WM Task | H | 0.17 | 0.01 | 19.72 | <.001 |
| | | | E | 0.11 | 0.01 | 13.24 | <.001 |
| | | | H>E | 0.06 | 0.01 | 5.46 | <.001 |
| right | hippocampus | Reading Language Task | S | 0.1 | 0.01 | 13.66 | <.001 |
| | | | N | 0.04 | 0.01 | 5.46 | <.001 |
| | | | S>N | 0.06 | 0.01 | 7.40 | <.001 |

| | | | | | | | |
|---|---|---|---|---|---|---|---|
| | | | S>H | 0.23 | 0.01 | 19.85 | <.001 |
| | | Auditory Language Task | I | 0.19 | 0.03 | 7.44 | <.001 |
| | | | D | 0.1 | 0.03 | 3.86 | <.001 |
| | | | I>D | 0.09 | 0.03 | 2.77 | .006 |
| | | | I>H | 0.31 | 0.03 | 11.97 | <.001 |
| | | Spatial WM Task | H | -0.13 | 0.01 | -11.90 | <.001 |
| | | | E | -0.05 | 0.01 | -4.30 | <.001 |
| | | | H>E | -0.08 | 0.01 | -6.07 | <.001 |
| right | amygdala | Reading Language Task | S | 0.14 | 0.01 | 13.52 | <.001 |
| | | | N | 0.06 | 0.01 | 5.97 | <.001 |
| | | | S>N | 0.08 | 0.01 | 7.22 | <.001 |
| | | | S>H | 0.27 | 0.02 | 17.71 | <.001 |
| | | Auditory Language Task | I | 0.44 | 0.03 | 13.21 | <.001 |
| | | | D | 0.21 | 0.03 | 6.35 | <.001 |
| | | | I>D | 0.23 | 0.04 | 5.38 | <.001 |
| | | | I>H | 0.57 | 0.03 | 16.49 | <.001 |
| | | Spatial WM Task | H | -0.13 | 0.01 | -8.98 | <.001 |
| | | | E | -0.06 | 0.01 | -4.34 | <.001 |
| | | | H>E | -0.07 | 0.02 | -3.82 | <.001 |
| right | accumbens | Reading Language Task | S | 0.02 | 0.01 | 2.33 | .020 |
| | | | N | 0.01 | 0.01 | 0.72 | .471 |
| | | | S>N | 0.02 | 0.01 | 1.46 | .145 |
| | | | S>H | -0.04 | 0.01 | -2.68 | .007 |
| | | Auditory Language Task | I | 0.02 | 0.03 | 0.52 | .600 |
| | | | D | 0.01 | 0.03 | 0.28 | .780 |
| | | | I>D | 0.01 | 0.04 | 0.19 | .850 |
| | | | I>H | -0.04 | 0.03 | -1.31 | .191 |
| | | Spatial WM Task | H | 0.06 | 0.01 | 4.44 | <.001 |
| | | | E | 0.06 | 0.01 | 4.46 | <.001 |
| | | | H>E | 0 | 0.02 | -0.01 | .989 |
