## supplemental table for "The extended language network: Language-responsive brain areas whose contributions to language remain to be discovered"

| BrainArea | Reference | Method | Modality | Task | Population |
|---|---|---|---|---|---|
| **left fronto-temporal cortex** | Goodglass, 1993 | review | | | lesions / aphasia |
| | Hickock & Poeppel, 2007 | review | | | |
| | Price, 2010 | review | | | |
| | Luria, 2011 | review | | | lesions / aphasia |
| | Friederici, 2012 | review | | | |
| | Price, 2012 | review | | | |
| | Hagoort, 2014 | review | | | |
| | Fridriksson et al., 2018 | review | | | lesions / aphasia |
| | Fedorenko et al., 2024 | review | | | |
| | Vigneau et al., 2006 | meta-analysis | | | |
| | Ferstl et al., 2008 | meta-analysis | | | |
| | Hodgson et al., 2021 | meta-analysis | | | |
| | Turker et al., 2023 | meta-analysis | | | |
| | Yang et al., 2024 | meta-analysis | | | deaf |
| | Kansaku et al., 2000 | fMRI | auditory comprehension | passive listening to stories | |
| | Pallier et al., 2003 | fMRI | auditory comprehension | sentences comprehension in native > unfamiliar language | |
| | Fedorenko et al., 2010 | fMRI | reading / auditory comprehension | passive reading / listening to sentences | |
| | Scott et al., 2017 | fMRI | auditory comprehension | passive listening to stories | |
| | Braga et al., 2020 | fMRI | resting state | | |
| | Lipkin et al., 2022 | fMRI | reading / auditory comprehension | passive reading / listening to sentences | |
| | Malik-Moraleda et al., 2022 | fMRI | auditory comprehension | passive listening to stories | |

| | | | | | |
|---|---|---|---|---|---|
| | Hu et al., 2022 | fMRI | language production | picture naming / picture description / word reading / sentence reading | |
| | Giglio et al., 2024 | fMRI | language production | naturalistic production / listening to naturalistic speech | |
| | Wolna et al., 2024 | fMRI | language production | picture naming | |
| | Rajimehr et al., 2024 | fMRI | naturalistic cognition | movie watching | |
| | Perani et al., 1996 | PET | auditory comprehension | passive story comprehension | |
| | Tzourio et al., 1998 | PET | auditory comprehension | passive story comprehension | |
| | Perani et al., 1998 | PET | auditory comprehension | passive listening to stories in native a foregin language | |
| | Papathanassiou et al., 2000 | PET | auditory comprehension / language production (covert) | passive story comprehension / verb generation (covert) | |
| | Alho et al., 2003 | PET | auditory comprehension | attentive listening to a story (left / right ear) | |
| | Crinion et al., 2003 | PET | auditory comprehension | passive listening to stories | |
| | Clark et al., 2005 | PET | langauge production / comprehension | | lesions / aphasia |
| | Friederici et al., 2000 | MEG | auditory comprehension | sentences / syntactic sentences / word lists / nonword lists | |
| | Broca, 1861 | case study | | | lesions / aphasia |
| | Wernicke, 1874 | case study | | | lesions / aphasia |
| **temporal pole** | Lambon Ralph et al., 2017 | review | | | |
| | Ferstl et al., 2007 | meta-analysis | | | |
| | Visser & Lambon-Ralph, 2011 | fMRI | auditory comprehension | semantic decision | |
| | Bédos Ulvin et al., 2017 | intracranial electrical stimulation | language production | object naming | epileptic patients |
| | Bedny et al., 2011 | fMRI | auditory comprehension in blind | sentence comprehension | blind |

| | | | | | |
|---|---|---|---|---|---|
| **ventral temporal areas** | Krauss et al., 1996 | intracranial electrical stimulation | language production / reading / auditory comprehension | object naming (BNT) / passage reading / spontaneous speech / responsive naming / single word reading | epileptic patients |
| | Snyder et al., 2023 | intracranial EEG / fMRI | language production | object naming (BNT) | epileptic patients |
| **medial parietal cortex** | Ferstl et al., 2007 | meta-analysis | | | |
| | Holland & Lambon Ralph, 2010 | rTMS | langauge production | generating verbs in past tense based on a present tense prompt | |
| **pre-SMA** | Hertrich et al., 2016 | review | | | |
| | Bohsali & Crosson, 2016 | review | | | |
| | Ardila, 2020 | review | | | lesions / aphasia |
| | Turkeltaub et al., 2002 | meta-analysis | | | |
| | Ferstl et al., 2007 | meta-analysis | | | |
| | Crosson et al., 2003 | fMRI | language production | word generation (based on thyming or semantic cues) | |
| **occipital: vOT / VWFA** | Reich et al., 2011 | fMRI | reading braille | word reading (Braille) | blind |
| | Kim et al., 2017 | fMRI | reading braille / auditory comprehension | word reading (Braille) / passive sentence comprehension | blind |
| **occipital: V1** | Seydell-Greenwald et al., 2023 | fMRI | auditory comprehension | listening to sentences with true / false judgement | |
| **occipital: vOT / VWFA** | Dikker et al., 2009 | MEG | reading | detecting syntactic violations | |
| **occipital: V1** | Dikker et al., 2010 | MEG | reading | detecting syntactic violations | |
| **hippocampus** | Brown-Schmidt, 2017, 2012 | review | | | |
| | Covington & Duff, 2016 | review | | | |
| | Tuckute, Lee, et al., in prep | fMRI | reading / auditory comprehension | passive reading / listening to sentences | |

| | | | | |
|---|---|---|---|---|
| | Wahl et al., 2008 | intracranial EEG / EEG | auditory comprehension | listening to syntactically correct vs. uncorrect sentences |
| | Piai et al., 2016 | intracranial EEG | auditory comprehension | listening to contextually constrained or unconstrained sentenes followed by picture naming |
| **cerebellum** | Leiner et al., 1993 | review | | |
| | Murdoch, 2010 | review | | |
| | Highnam & Bleile, 2011 | review | | |
| | Mariën et al., 2014 | review | | |
| | Ackerman & Brendel, 2016 | review | | |
| | Van Dun et al., 2016 | review | | |
| | LeBel & D'Mello, 2023 | review | | |
| | Casto et al., 2025 | fMRI | auditory comprehension / reading / language production | passive reading / listening to sentences |
| **thalamus** | Klostermann & Ehlen, 2013 | review | | |
| | Booth et al., 2007 | fMRI | reading / rhyming task | single word reading with rhyming judgement |
| **basal ganglia** | Kotz et al., 2009 | review | | |
| | Ackerman & Brendel, 2016 | review | | |
| | Crosson et al., 2003 | fMRI | language production | word generation (based on thyming or semantic cues) |
| | Thibault et al., 2021 | fMRI | reading | reading sentences with true vs. false judgement |
